# Proteomes Reveal Metabolic Capabilities of *Yarrowia lipolytica* for Biological Upcycling of Polyethylene into High-Value Chemicals

**DOI:** 10.1101/2023.04.17.537167

**Authors:** Caleb Walker, Max Mortensen, Bindica Poudel, Christopher Cotter, Ikenna Okekeogbu, Seunghyun Ryu, Bamin Khomami, Richard J. Giannone, Siris Laursen, Cong T. Trinh

## Abstract

Polyolefins derived from plastic wastes are recalcitrant for biological upcycling. However, chemical depolymerization of polyolefins can generate depolymerized plastic (DP) oil comprising of a complex mixture of saturated, unsaturated, even and odd hydrocarbons suitable for biological conversion. While DP oil contains a rich carbon and energy source, it is inhibitory to cells. Understanding and harnessing robust metabolic capabilities of microorganisms to upcycle the hydrocarbons in DP oil, both naturally and unnaturally occurring, into high-value chemicals are limited. Here, we discovered that an oleaginous yeast *Yarrowia lipolytica* undergoing short-term adaptation to DP oil robustly utilized a wide range of hydrocarbons for cell growth and production of citric acid and neutral lipids. When growing on hydrocarbons, *Y. lipolytica* partitioned into planktonic and oil-bound cells with each exhibiting distinct proteomes and amino acid distributions invested into establishing these proteomes. Significant proteome reallocation towards energy and lipid metabolism, belonging to two of the 23 KOG (Eukaryotic Orthologous Groups) classes C and I, enabled robust growth of *Y. lipolytica* on hydrocarbons, with n-hexadecane as the preferential substrate. This investment was even higher for growth on DP oil where both the KOG classes C and I were the top two, and many associated proteins and pathways were expressed and upregulated including the hydrocarbon degradation pathway, Krebs cycle, glyoxylate shunt and, unexpectedly, propionate metabolism. However, a reduction in proteome allocation for protein biosynthesis, at the expense of the observed increase towards energy and lipid metabolisms, might have caused the inhibitory effect of DP oil on cell growth.

**MPORTANCE:** Sustainable processes for biological upcycling plastic wastes in a circular bioeconomy are needed to promote decarbonization and reduce environmental pollution due to increased plastic consumption, incineration, and landfill storage. Strain characterization and proteomic analysis revealed the robust metabolic capabilities of *Y. lipolytica* to upcycle polyethylene into high-value chemicals. Significant proteome reallocation towards energy and lipid metabolisms was required for robust growth on hydrocarbons with n-hexadecane as the preferential substrate. However, an apparent over-investment in these same categories to utilize complex DP oil came at the expense of protein biosynthesis, limiting cell growth. Taken together, this study elucidates how *Y. lipolytica* activates its metabolism to utilize DP oil and establishes *Y. lipolytica* as a promising host for the upcycling of plastic wastes.

## INTRODUCTION

Plastics are hydrocarbons that are mainly derived from petroleum (1). Its high demand and production with limited recyclability generate plastic wastes that have been polluting our ecosystems for decades, negatively impacting both wildlife and human health (1, 2). Yet, sustainable processes to recycle plastic wastes remain elusive since current recycling technologies produce polymers with lower quality and higher cost (3). One solution to improve the sustainability and cost competitiveness of the recycling process is to upcycle plastic wastes into high-value chemicals (4). Due to the abundance of hydrocarbons in plastic wastes such as polyolefins (e.g., polyethylene, polypropylene), efficient biological upcycling of these wastes into high-value chemicals at standard conditions, such as room temperature and atmospheric pressure, is both disruptive and transformative (5).

In nature, certain microorganisms have been discovered for their unique metabolic capability to degrade plastic (6, 7). However, direct microbial depolymerization of plastic wastes, especially those derived from complex and recalcitrant polyolefins, is inefficient due to slow degradation rates that take years to complete (7-9). One strategy to overcome this limitation is to first depolymerize plastic wastes in a thermochemical pretreatment step to generate low molecular weight depolymerized plastic (DP) intermediates (e.g., C11-C28 hydrocarbons in a form of oil and wax) (10) that microorganisms can rapidly and effectively utilize for biosynthesis of high-value chemicals with high selectivity and ease of separation in the biological upcycling step. This strategy resembles an integrated biorefinery for conversion of lignocellulosic biomass to biofuels and biobased products where chemical pretreatment is first performed to reduce biomass recalcitrance followed by saccharification and fermentation to produce target products (11).

For thermochemical pretreatment, catalytic cleavage of polyolefins to C11-C28 hydrocarbon intermediates that microorganisms can utilize is more advantageous than less controlled methods such as non-catalytic pyrolysis or direct functionalization (12-15). The catalytic approach enables the reaction to be accelerated at lower temperatures than pyrolysis and can provide far greater control over product distributions (16-18). As polyolefins are large, saturated hydrocarbons, solid acid catalysts are commonly employed due to the well-known surface chemistry of acidic sites in hydrocarbon activation, cleavage, and isomerization (19-24). The elementary mechanism for solid-acid-catalyzed saturated hydrocarbon cleavage proceeds through dehydrogenation, C-C bond cleavage, and either desorption or re-hydrogenation to produce unsaturated or saturated products in a range of C11-C28 hydrocarbon intermediates suitable for biological conversion (19, 20, 25). Depending on reaction temperature, hydrogenation kinetics over solid acid catalysts can be kinetically slow, thus metal co-catalysts might need to be employed to improve the saturation of products (26-29). For biological upcycling, the resulting hydrocarbon intermediates contain rich carbon and electron sources that enable microorganisms to grow and produce high-value chemicals. However, mixed and complex hydrocarbons with saturated, unsaturated, linear, branched, even and/or odd carbon chains can be inhibitory and non-degradable to microorganisms since they have not yet adapted to utilize these hydrocarbons. Therefore, biological upcycling of depolymerized plastics to produce high-value chemicals requires efficient and robust microorganisms.

One promising microbial host that can be repurposed for biological upcycling of hydrocarbon intermediates is the generally-regarded-as-safe (GRAS certified) oleaginous yeast *Yarrowia lipolytica* (30, 31). This organism has broad industrial use with the unique metabolic capability to efficiently produce high-value chemicals (e.g., organic acids (32-35), erythritol (36), lipid-derived products (37-41)), has tractable genetics with facile genetic manipulation (42), and demonstrates natural robustness, being able to thrive in a broad range of pH (43), high salinity (44), organic solvents such as ionic liquids (45-47), and inhibitory biomass hydrolysates (48, 49). Since *Y. lipolytica* was isolated from fat-rich environments such as cheese and sausage (50), it is also known to naturally assimilate C10-C18 n-alkanes (or paraffins), with n-hexadecane (C16) as the most preferable substrate (31, 51, 52).

The n-alkane degradation process is complex*. Y. lipolytica* first secretes an emulsifier (e.g., liposan) to sequester alkanes for targeted uptake through cell adhesion and cellular protrusions (53-55). Intracellular n-alkane degradation involves multiple enzymatic steps across several subcellular compartments including the cytosol, endoplasmic reticulum (ER), peroxisome, and mitochondria (56). In the ER, n-alkanes are oxidized to fatty alcohols by cytochrome P450s (57, 58), then to fatty aldehydes by fatty alcohol dehydrogenases and/or fatty alcohol oxidases (59), and finally to fatty acids by fatty aldehyde dehydrogenases (60). Once transported into the peroxisome, fatty acids are converted to fatty acyl-CoAs by ligases and degraded through the beta-oxidation cycle (56, 61). The terminal metabolite of the n-alkane degradation in the peroxisome is acetyl-CoA, a building block for cell growth and product synthesis. In mitochondria, acetyl-CoA enters the Krebs cycle, generating energy and producing dicarboxylic acids (e.g., citric acid, alpha-ketoglutaric acid, succinic acid). However, in the cytosol it is directed toward lipid biosynthesis. Even though some metabolic processes involved in *Y. lipolytica*’ s n-alkane degradation have been characterized, it is still largely unknown how *Y. lipolytica* utilizes DP intermediates derived from plastic wastes (e.g., polyolefins) that contain a complex mixture of hydrocarbons.

In this study, we generated depolymerized polyethylene by thermocatalytic pretreatment and investigated the robustness of *Y. lipolytica* for biological upcycling of the mixed and complex (C11-C28) hydrocarbons. Through a short-term adaptation, we isolated an adapted *Y. lipolytica* and characterized its ability to utilize the inhibitory DP intermediates for growth and production of citric acid and lipids. We elucidated the robust cellular processes of *Y. lipolytica* for utilizing these complex hydrocarbons and quantitatively determined how *Y. lipolytica* allocated its proteome for cell growth and product formation.

## RESULTS

### Catalytic depolymerization of low linear density polyethylene (LLDPE) with SiC catalyst generated DP intermediates suitable for microbial conversion

To generate depolymerized plastic intermediates that are compatible with biological upcycling by *Y. lipolytica*, we targeted the production of saturated, linear C11-C18 hydrocarbons as DP oil. As a proof-of-study, we investigated catalytic depolymerization of LLDPE as a model polyolefin waste in a static bed reactor with a condensation flow under an inert gas of CO_2_ (Figure 1A) using an unmodified SiC catalyst (Figure 1B). The surface composition of SiC is a suboxide of SiO, which presents weak acid sites to drive polyolefin cleavage. Since the polymer itself autogenously provides atomic hydrogens for the hydrogenation of surface-bound cleavage reaction intermediates, no external hydrogen source was added. A static reaction bed was dewatered, and the reaction proceeded at 400°C for 8 hours. To maximize favorable oil production, we manipulated the catalyst to polymer ratios of 0, 0.25, 1.25, and 2.5.

**Figure 1.**
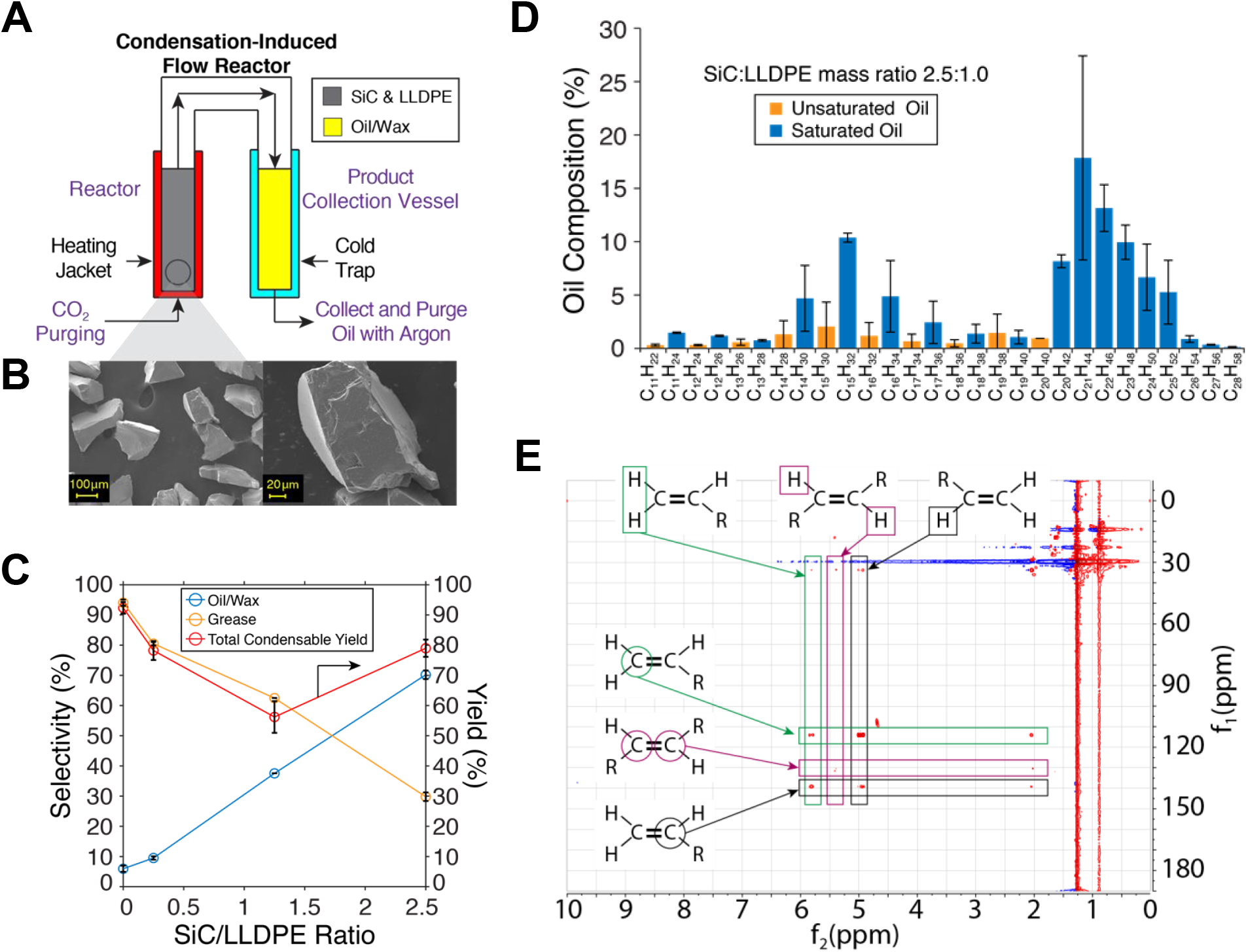
(**A**) Scheme of a condensation-induced flow reactor built for thermocatalytic depolymerization of plastics. (**B**) SEM micrograph of the SiC catalyst. (**C**) Selectivities and yields for LLDPE depolymerization in a semi-batch reactor with a range of SiC-to-LLDPE ratios. Reactions were performed at 400°C for 8 h under an inert gas of CO_2_. (**D**) Composition of DP oil/wax analyzed by GCMS. (**E**) Analysis of (un)Saturation of DP oil/wax by 1H-13C HMBC NMR.

Thermocatalytic depolymerization of LLDPE generated four types of product streams, including grease, oil, wax, and low molecular weight hydrocarbon off-gases. The reaction without the SiC catalyst resulted in an overall condensable yield of 91% with a grease selectivity of 95% and an oil/wax selectivity of 5% (Figure 1C). Introduction of SiC at a SiC:LLDPE ratio of 0.25 produced a higher oil/wax selectivity of 18% but the total condensable yield decreased to 81%. With a SiC:LLDPE ratio of 1.25, the oil/wax selectivity increased to 41% with a drastic decrease in total condensable yield down to 55%. With a SiC:LLDPE ratio of 2.5, the oil/wax selectivity further improved to 72% and the total condensable yield increased to 78%. These results illustrate the role of thermal radicals in the formation of the grease product at 400°C when no catalyst was present. Access to the catalytic surface chemistry of SiC was demonstrated by a monotonic increase of wax and oil yield as catalyst to polymer ratio increased. Unlike the oil/wax yields, the total condensable yields did not track monotonically with the catalyst-to-polymer ratios, which underlines the significant role of mass transport limitation in the reaction.

Through the GCMS and 1H-13C HMBC NMR analyses, we found that the oil/wax product consisted of mostly saturated linear hydrocarbon chains of size C_11_-C_28_ and exhibited a bi-modal product distribution with modes centered at C_15_ and C_21_ (Figures 1D, 1E). Estimated saturation by GCMS was found to be 96% with only 4% unsaturated hydrocarbons. Characterization of the oil by 1H–13C HMBC NMR (Figure 1E) indicated that end-chain unsaturation dominated the distribution of unsaturated products. Gel permeation chromatography of the grease product indicated an average molecular weight of 69,630 g/mol. The results suggest that polymer chain dynamics could play a significant role in determining the molecular distribution of hydrocarbons within the oil product and may provide an avenue for improved catalytic performance if appropriate surface chemistry could be isolated. The bi-modal molecular distribution observed implies that mid-chain stress dynamics contributed to the catalytic cleavage. If mid-chain stress were not playing a role, there would be no selectivity for C-C cleavage beyond that of end-chain and inner-chain bond with the former known to be more energetically demanding and a flatter molecular distribution would be expected (62-64).

Overall, the thermocatalytic depolymerization of LLDPE in a condensation-induced flow reactor using SiC catalyst with SiC:LLDPE of 2.5:1.0 generated good yields and selectivities of oil/wax product streams that can be suitable for downstream bioprocessing. The next step was to characterize and elucidate the compatibility of *Y. lipolytica* for biological upcycling of DP oil.

### *Y. lipolytica* used DP oil as carbon and energy sources for growth

#### Y. lipolytica was capable of growing on DP oil

Initial characterization of the parent *Y. lipolytica* Po1f showed cell growth (i.e., turbidity) in culture tubes containing 5% (v/v) DP oil as a sole carbon source and in culture tubes containing a mixture of 5% (v/v) DP oil and 5 g/L glucose after 2 days of cultivation (Figure 2A). To confirm cell growth on DP oil, we repeated this experiment using baffled flasks to provide better aeration for cell growth. We found that 2% (v/v) DP oil as a sole carbon source supported growth of *Y. lipolytica* cells (Figure 2B). However, cell growth was not detected using 5% or 10% (v/v) DP oil. Notably, maximum cell density (i.e., OD_600nm_) in a mixture of glucose and DP oil was reduced by ∼53% in comparison to cell growth in glucose alone (Figure 2B). These results indicate DP oil is inhibitory to *Y. lipolytica*.

**Figure 2.**
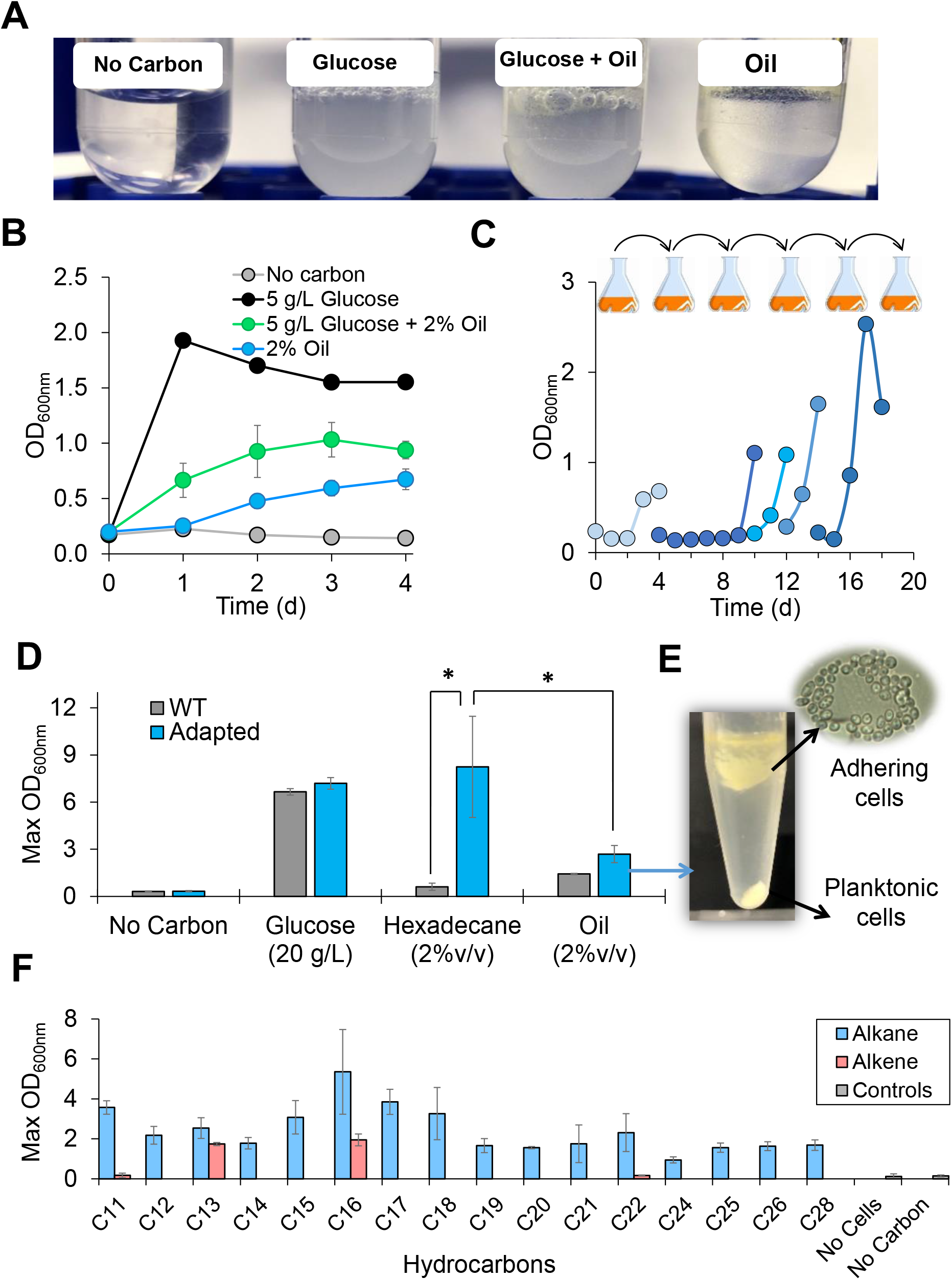
Comprehensive growth characterization of *Y. lipolytica* in hydrocarbons. (**A-B**) Growth of the parent *Y. lipolytica* Po1f in (**A**) culture tubes and (**B**) baffled flasks. (**C**) Short-term adaptation of *Y. lipolytica* in DP oil to generate adapted strain. (**D**) Growth comparison of the parent and adapted *Y. lipolytica* in glucose, n-hexadecane, and DP oil. Symbol: *p-value < 0.001 using one-way analysis of variance Holm-Sidak method. (**E**) Image of adapted *Y. lipolytica* cultured in DP oil after centrifugation and microscopic image of adhering (or bound) cells. **(F)** Growth of DP-adapted *Y. lipolytica* on individual n-alkanes (blue) and 1-alkenes (red).

#### Short-term adaptation of *Y. lipolytica* significantly enhanced cell growth on DP oil

To improve cell growth on DP oil, we employed a short-term adaptation experiment by subjecting wildtype Y. lipolytica to 5 successive transfers in 2% (v/v) oil (Figure 2C). Short-term adaptation (∼10 generations) improved growth performance in DP oil, enabling the adapted strain (maximum OD600nm of 2.69 ± 0.55) to increase cell density ∼1.9-fold as compared to the parent strain (maximum OD600nm of 1.43 ± 0.04) using 2% (v/v) DP oil as a sole carbon source (Figure 2D). Compared to DP oil, the adapted strain demonstrated superior growth (maximum OD600nm of 8.24 ± 0.55) on the model substrate n-hexadecane, a component of DP oil, by 3.1-fold (Figure 2D). In addition to observing planktonic cells in the cultures, we found that the adapted strain exhibited cellular adhesion to the hydrophobic layer of both DP oil and n-hexadecane even after centrifugation (Figure 2E).

#### Adapted Y. lipolytica exhibited improved growth on a broad range of saturated n-alkanes and some select 1-alkenes present in DP oil/wax

While *Y. lipolytica* is known to grow on C11- C18 paraffins (oil form) (31, 51, 52), it has not yet been shown to utilize longer n-alkanes (wax form) or 1-alkenes that are also present in DP intermediates. To better understand how *Y. lipolytica* could utilize these types of DP intermediates (oil/wax) (Figure 1D), we next characterized and compared cell growth of adapted *Y. lipolytica* across a variety of individual hydrocarbons. Interestingly, all (C11-C28) n-alkanes supported growth of DP oil-adapted *Y. lipolytica* as the sole carbon sources (Figure S1). Notably, cell growth on (C16) n-hexadecane outperformed all other hydrocarbons tested (Figure 2F). However, only medium carbon chain length alkenes (i.e., 1-C13 and 1-C16) supported cell growth, whereas cells failed to grow with short (i.e., 1-C11) or long (i.e., 1-C22) alkenes as the sole carbon sources (Figure 2F, S2). Taken together, *Y. lipolytica* has the metabolic capability to utilize the complex mixture of alkanes and alkenes in the DP oil derived from polyethylene. Medium-chain saturated hydrocarbons (e.g., C11-C18) best supported the growth of *Y. lipolytica*.

#### Instability of DP oil in long-term storage became inhibitory to *Y. lipolytica*

During our experiments using DP oil, we observed reduced cell growth from DP oil stored at room temperature (RT) for a long period of time (∼2-3 months). To test whether the instability of DP oil could increase its toxicity to cells, we partitioned the fresh DP oil into two identical batches for storage at two conditions, i.e., one at RT and the other at −20℃. Growth characterization of adapted *Y. lipolytica* at 3 and 6 weeks of storage showed no statistical differences in cell growth between the two storage conditions, albeit RT-stored oil resulted in greater deviation of growth between replicates (Figure 3A). After 9 weeks, the deviation in cell growth between replicates became prominent for RT-stored oil (Figure 3A). Finally, after 14 weeks of storage, the adapted strain was unable to grow in the RT-stored oil while growth was not affected by oil stored at −20℃ (Figure 3A). To better understand the unstable components of the DP oil, we subjected oil from both storage conditions at 14 weeks to 2D NMR and FTIR analyses. While NMR revealed no distinguishable differences between the two storage conditions (Figure S1), FTIR revealed one band in the RT-DP oil scan at 1707cm^-1^ that deviated from the −20°C stored oil which indicates carbonyl groups from either ketones, esters, or acids are present (Figure 3B). The bands associated with esters and acids were not present in the RT-DP oil scan, indicating that the accumulation of ketones in RT-stored DP oil likely caused growth inhibition (Figure 3C).

**Figure 3.**
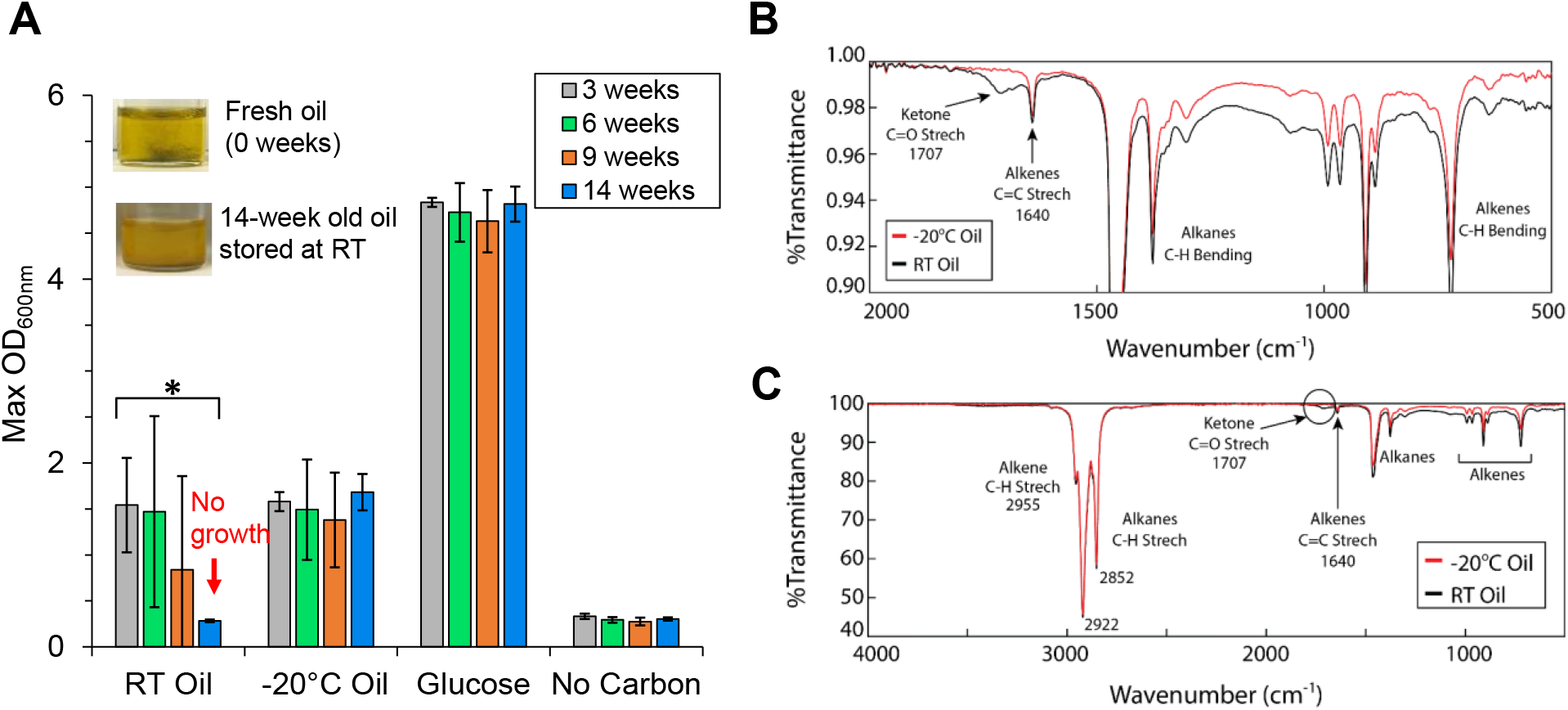
Effect of DP oil stability on growth of adapted *Y. lipolytica*. (**A**) Growth of adapted *Y. lipolytica* on DP oil stored at RT or −20°C after 3, 6, 9 and 14 weeks. (**B-C**) FTIR analysis of 14- week DP oil stored at RT and −20°C **(B)** with a zoom-in wavelength range of 500-2000 cm^-1^ and **(C)** with a zoom-in wavelength range of 500-4000 cm^-1^. *p-value < 0.001 using one-way analysis of variance Holm-Sidak method.

### *Y. lipolytica* upcycled DP oil to citric acid and lipids

Next, we investigated whether *Y. lipolytica* could upcycle DP oil into higher-value chemicals, specifically triacylglycerol lipids and citric acid, using nitrogen limited (C:N = 100) media. We also used glucose and n-hexadecane as controls. Using 2% (v/v) DP oil, adapted *Y. lipolytica* achieved a maximum cell density of 3.34 ± 0.32 OD_600nm_ (Figure 4A) while producing 2.33 ± 0.12 g/L citric acid (Figure 4B) and accumulating 10.09 ± 0.42% lipid in cell mass (Figure 4C). However, maximum cell density (11.74 ± 0.36 OD600_nm_) and citric acid production (14.6 ± 0.77 g/L citric acid) were significantly higher in 2% (v/v) n-hexadecane than in the DP oil (Figure 4A, 4B). Both n-hexadecane and DP oil carbon sources exhibited similar lipid levels by day 4 (Figure 4C). Cells grown in 20 g/L glucose showed the lowest lipid level (6.93 ± 1.17% lipid in cell mass) with moderate citric acid titer (4.28 ± 0.37 g/L citric acid) and intermediate cell growth (7.34 ± 0.14 OD600nm) between the three carbon sources (Figure 4A, 4C). Interestingly, both n-hexadecane and DP oil GCMS profiles showed complete consumption despite the differences in cell growth and citric acid production (Figure 4D). Cultivation with only n-hexadecane enabled superior growth of *Y. lipolytica* and its production of citric acids and lipids, especially relative to the DP oil. While adapted *Y. lipolytica* could consume DP oil, its growth was limited because the mixed and complex composition of the oil might have limited cellular metabolism impacting substrate assimilation and hence growth and product formation. To understand these differences, we next employed proteomics to elucidate how *Y. lipolytica* utilized hydrophobic hydrocarbons.

**Figure 4.**
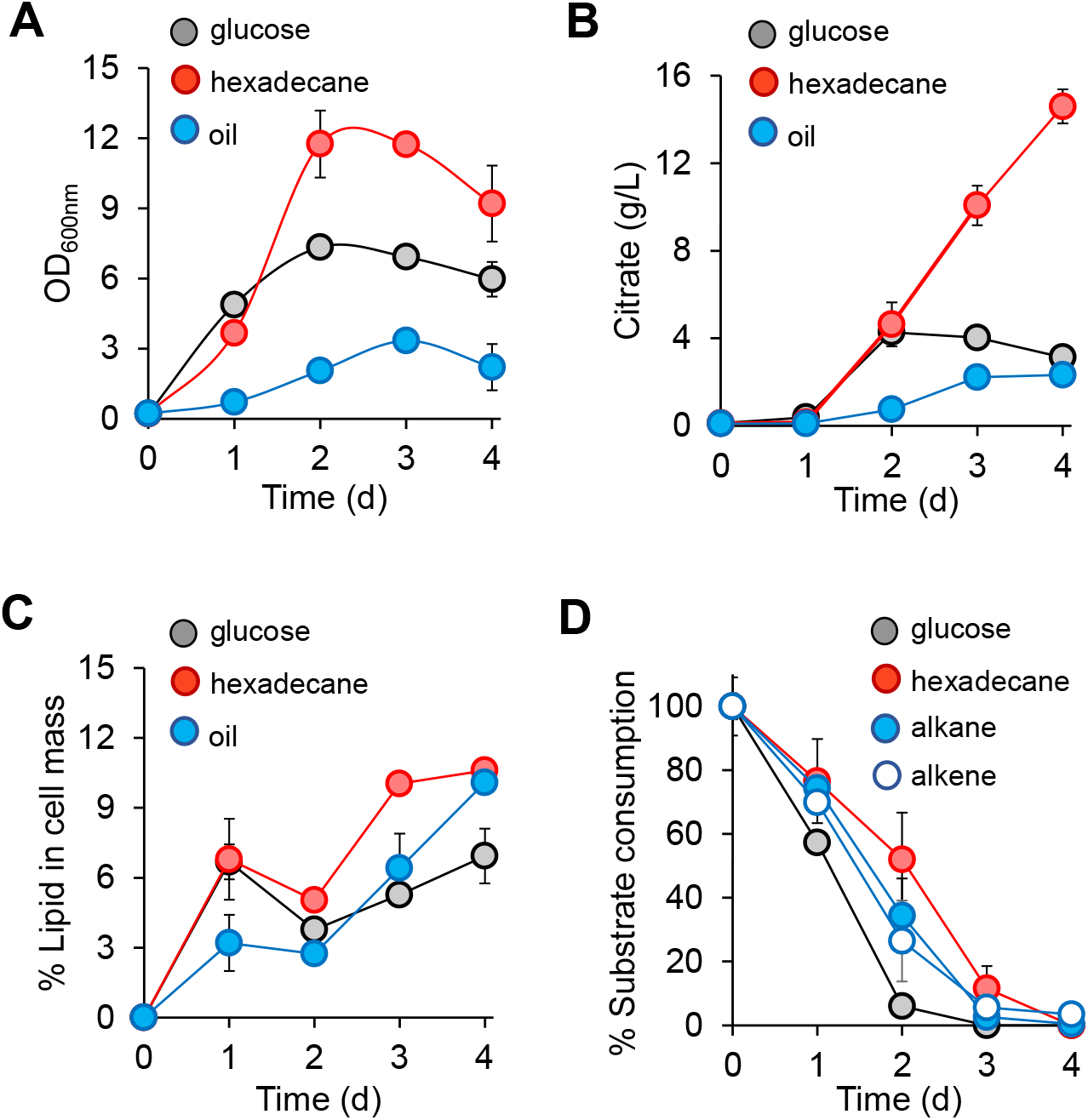
Biological upcycling of DP oil by adapted *Y. lipolytica* for production of citric acid and neutral lipids. (**A**) Growth kinetics of adapted *Y. lipolytica* on DP oil and control substrates (glucose and n-hexadecane), (**B**) Citric acid production, (**C**) Lipid accumulation, and (**D**) substrate consumption profiles.

### Proteomes revealed robust metabolic capability of *Y. lipolytica* for utilizing n-hexadecane

#### Planktonic and bound cells exhibited distinct proteomes

To elucidate the metabolic capability of the adapted *Y. lipolytica* for oil upcycling, we first compared proteomes of i) planktonic cells (GP24) growing on glucose (control), ii) planktonic cells (HP24) growing on n-hexadecane, and iii) bound (or oil-adhering) cells (HB24) growing on n-hexadecane during the exponential growth phase after day 1 (Figures 5A, 5B). Principal component analysis (PCA) showed that GP24, HP24, and HB24 cells exhibited distinct proteomes (Figure 5C). Both HP24 and HB24 cells shared 211 commonly upregulated and 82 commonly downregulated proteins as compared to GP24 cells (Figure 5D). The annotated, differentially expressed proteins represented all of the 23 KOG classes but had distinct up/down regulation patterns based on the growth conditions (Figures 5E, 5F, 5G). The HP24 cells had 415 upregulated proteins (log2 > 1, p-value <0.05, Student’s T-test) and 118 downregulated proteins (log2 < −1, p-value <0.05, Student’s T-test) belonging to numerous KOG classes (Figures 5D, S3A, Data Set S1). However, HB24 cells upregulated 241 proteins and downregulated 242 proteins (Figures 5F, S3B, Data Set S2). Pairwise comparison between planktonic HP24 and bound HB24 cells showed that HP24 cells had 388 upregulated proteins and 71 downregulated proteins (Figures 5G, S3C, Data Set S3). Hierarchical clustering across all of the samples also confirmed distinct differences between the proteomes of planktonic and bound cells. (Figure S3D).

**Figure 5:**
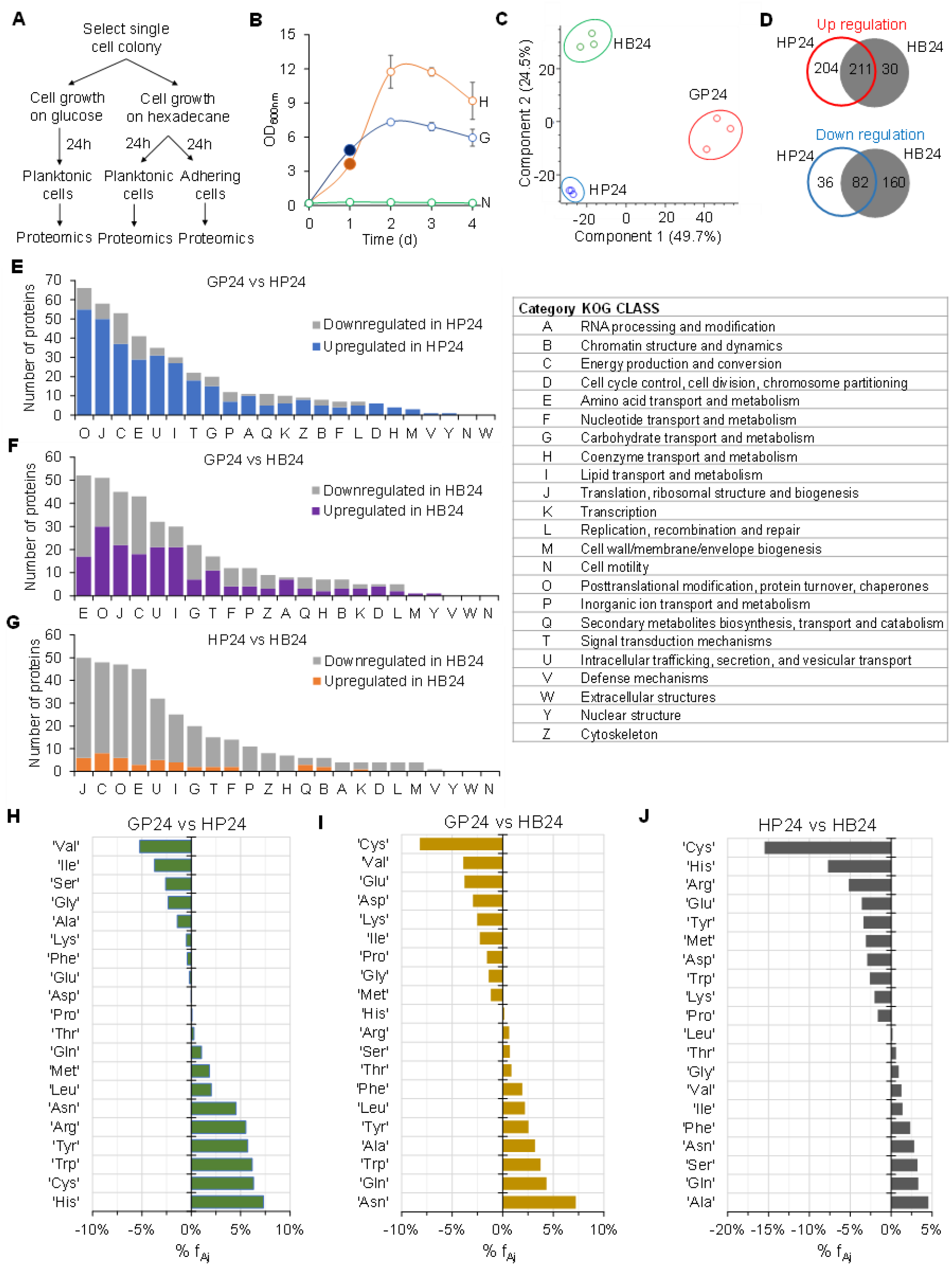
Distinct proteomes of *Y. lipolytica* cells growing on glucose and n-hexadecane. **(A)** Experimental design. **(B)** Growth and cell sample collection for proteomic analysis. Filled circle at day 1 indicates that samples were collected for proteomic analysis. Abbreviations: H, hexadecane; G, glucose; N, no carbon. **(C)** Principal component analysis of proteomes of GP24, HP24, and HB24 cells. **(D)** Proteome comparison between GP24, HP24, and HB24 cells. Regulated proteins between two biological conditions are defined with p-values less than 0.05 and log_2_ fold changes that are either greater than 1 (upregulated proteins) or less than −1 (downregulated proteins). (**E-G**) Number of regulated proteins between (**E**) HP24 and GP24, (**F**) HB24 and GP24, and (**G**) HB24 and HP24. **(H-J)** Percent change of mass fraction of amino acids in the measured proteomes between (**H**) HP24 and GP24, (**I**) HB24 and GP24, and (**J**) HP24 and HB24.

#### Investment of amino acids in proteomes of adapted *Y. lipolytica* varied for growth on different substrates

Analyzing mass fractions of amino acids in the proteomes further demonstrates the global reprogramming of cellular metabolism of adapted *Y. lipolytica* growing on different substrates. As compared to the GP24 cells, the mass fraction of amino acids in the measured proteome of HP24 cells increased by 5% or more for Arg (+5.53%), Tyr (+5.72%), Trp (+6.17%), Cys (+6.31%), and His (+7.30%) and decreased 5% or more for Val (−5.23%) (Figure 5H). In contrast, the mass fraction of amino acids in the measured proteome of HB24 cells decreased by 5% or more only for Cys (−8.20%) while it increased 5% or more for Asn (+7.23%) relative to GP24 (Figure 5I). When comparing the planktonic (HP24) and bound (HB24) cells growing on oil, the mass fraction of amino acids in the measured proteome in HB24 cells decreased by 5% or more for Arg (−5.17%), His (−7.72%) and Cys (−15.49%) (Figure 5J). The high percentage changes in specific amino acids clearly show the effect on cellular resource utilization when *Y. lipolytica* redistributed its proteome to compensate for different environmental conditions. Modulating the availability of these amino acids via either *in vivo* biosynthesis or external supplementation can improve the fitness of *Y. lipolytica* for growth on hydrocarbons.

#### Proteome reallocation of energy and lipid metabolism enabled adapted *Y. lipolytica* to grow efficiently on n-hexadecane

To understand how GP24, HP24, and HB24 cells invested cellular resources for growth on different carbon sources, we analyzed the mass fractions of pathway proteins classified in 23 KOG classes. Regardless of growth conditions, we found that the top 7 KOG classes ─ including i) Energy production and conversion (C), ii) Translation, ribosomal structure and biogenesis (J), iii) Carbohydrate transport and metabolism (G), iv) Lipid transport and metabolism (I), v) Amino acid transport and metabolism (E), vi) Posttranslational modification, protein turnover, chaperones (O), and vii) Inorganic ion transport and metabolism (P), ─ made up about 70-75% of the total measured proteome abundance in adapted *Y. lipolytica* (Figure 6B, S4). Among the KOG classes, proteome allocation was most invested in energy metabolism (C) and protein biosynthesis (J, E, O).

**Figure 6:**
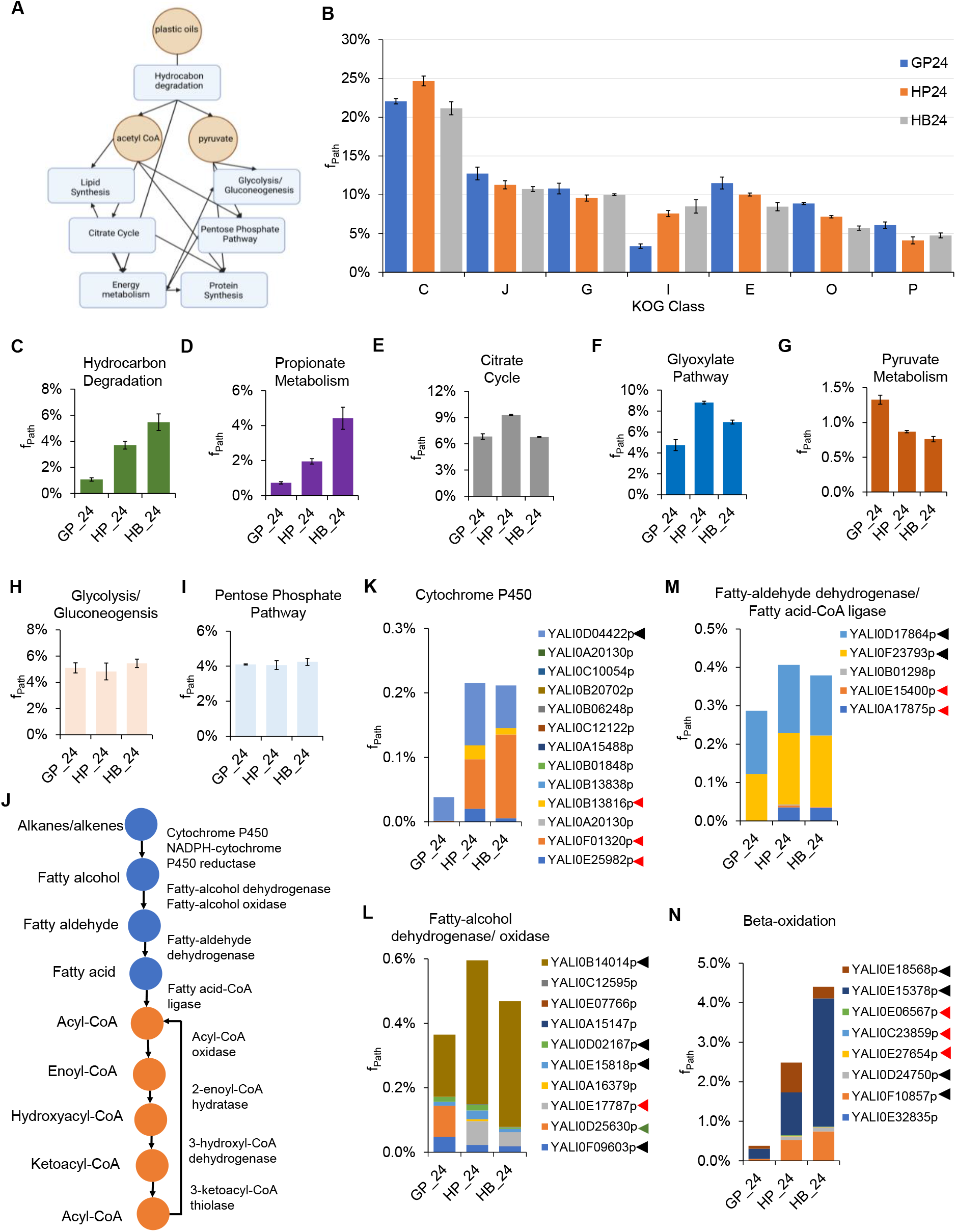
Analysis of proteome reallocation enabling robust growth of *Y. lipolytica* on hydrocarbons. (**A**) Key metabolic pathways for catabolic and anabolic metabolism of hydrocarbon utilization. **(B-K)** Comparison of mass fractions of proteomes invested in **(B)** the top 7 KOG classes, **(C)** hydrocarbon degradation pathway, (**D**) propionate metabolism, **(E)** citrate cycle, **(F)** glyoxylate pathway, **(G)** pyruvate metabolism, **(H)** glycolysis/gluconeogenesis, and **(I)** pentose phosphate pathway for growth on glucose, n-hexadecane, and DP oil. (**J**) Metabolic pathway for hydrocarbon degradation. (**K-N**) Mass fractions of **(K)** cytochrome P450s, **(L)** fatty alcohol dehydrogenases/oxidases, **(M)** fatty aldehyde dehydrogenases/fatty acid-CoA ligases, and **(N)** beta-oxidation proteins in the measured proteomes. Symbols: Black arrows: proteins present in GP24, HP24, and HB24; Red arrows: proteins present in HP24 and HB24 only; and green arrow: proteins present in GP24 only.

When growing on n-hexadecane, HP24 cells increased proteome allocation in the lipid transportation and metabolism KOG class (I) by 124% and energy production and conversion KOG class (C) by 12% relative to GP24 cells. This resource allocation came at the expense of other KOG classes (Figure 6B). For the KOG class I, there was a total of 27 upregulated proteins (log2 > 1, p-value <0.05, Student’s T-test) (Data Set S1) that are involved in hydrocarbon transport and degradation. For the top 5 upregulated proteins, YarlipO1F2_226951 (Q6C4Y2, YALI0E22781p, oxysterol-binding protein) was upregulated by 68-fold (p-value = 0), YarlipO1F2_193467 (Q6CHL1, YALI0A07733p, enoyl-CoA isomerase) by 60-fold (p-value = 0.0011), YarlipO1F2_230457 (F2Z6J3, YALI0F01320p, cytochrome P450) by 55-fold (p-value = 0.0038), YarlipO1F2_232715 (O74936, YALI0D02475p, acyl-CoA oxidase by 48-fold (p-value = 0.0012), and YarlipO1F2_277873 (Q6CGL4, YALI0A18337p, lysophospholipase) by 37-fold (p-value = 0).

For the KOG class C, 37 proteins were upregulated (log2 > 1, p-value <0.05, Student’s T-test) (Data Set S1); these highly upregulated proteins belong to the electron transport and Krebs cycle. Among these, the top 5 upregulated proteins include: YarlipO1F2_232437 (Q6C877, YALI0D22022p, mitochondrial F1F0-ATP synthase) upregulated by 376-fold (p-value = 0.0041), YarlipO1F2_208678 (Q6C4G4, YALI0E27005p, pyruvate dehydrogenase) by 60-fold (p-value = 0.0183), YarlipO1F2_233609 (Q6CCU5, YALI0C06446p, proteins containing the FAD binding domain) by 55-fold (p-value = 0.0129), YarlipO1F2_238445 (Q6CGH0, YALI0A19448p, aldehyde dehydrogenase) by 48-fold (p-value = 0.0044), and YarlipO1F2_86039 (Q6C7X2, YALI0D24629p, and acyl carrier protein by 38-fold (p-value = 0.0057). For HB24 cells, only the lipid transportation and metabolism (KOG class I) increased up to 152% relative to GP24 cells (Figure 6B). There was a total of 21 upregulated proteins; like HP24 cells, the highly upregulated proteins of HB24 cells participated in the hydrocarbon transport and degradation (Data Set S2).

For the pairwise comparison between the planktonic (HP24) and bound (HB24) cells growing on n-hexadecane, we found that HB24 cells increased proteome allocation to three KOG classes: 5% in the carbohydrate transport and metabolism (KOG class G), 12% in the lipid transport and metabolism (KOG class I) and 15% in the inorganic ion transport and metabolism (KOG class P) (Figure 6B). Since HB24 cells aggregated upon and sequestered n-hexadecane directly, higher proteome investments were made into these KOG classes, which are likely beneficial for growth on hydrophobic substrates.

#### Hexadecane induced a proteome reallocation toward hydrocarbon degradation, Krebs cycle, glyoxylate cycle, and propionate metabolism

To support cell growth and maintenance, living cells are required to synthesize the key precursor metabolites, such as acetyl CoA and pyruvate, which are important to make cellular building blocks (Figure 6A). Further examination of the perturbed KOG classes using annotated KEGG pathways/metabolism could provide a more granular view of how *Y. lipolytica* reprogrammed their metabolism to utilize n-hexadecane. To grow on n-hexadecane, *Y. lipolytica* reallocated a significant mass fraction of its proteome into the hydrocarbon degradation pathway (Figure 6C). Relative to HP24, HB24 cells invested their proteome resources towards hydrocarbon degradation by 1.5-fold and compared to GP24 by 5.1-fold. This result showed that not only was a greater proteomic investment made in hydrocarbon degradation, but also that more proteins were present and upregulated.

The metabolic pathway for hydrocarbon degradation is shown in Figure 6J. *Y. lipolytica* has a set of 12 cytochrome P450s that enable it to grow on a wide range of paraffins by oxidizing them into fatty alcohols (57, 58). We found that *Y. lipolytica* utilized a subset of 4 proteins including YALI0E25982p, YALI0F01320p, YALI0B13816p, and YALI0D04422 where the first three proteins were unique for growth on n-hexadecane (Figure 6K). Among the set of 9 fatty alcohol dehydrogenases and 1 alcohol oxidase used to synthesize fatty aldehydes, four of these proteins including YALI0F09603p, YALI0E15818p, YALI0D02167p, and YALI0B14014p were utilized for both growth on both glucose and n-hexadecane; and YALI0D25630p was unique to growth on glucose while YALI0E17787p was unique to growth on n-hexadecane (Figure 6L). To convert fatty aldehydes to fatty acids, *Y. lipolytica* utilized 3 (YALI0A17875p, YALI0E15400p, and YALI0F23793p) out of 4 available proteins where YALI0A17875p and YALI0E15400p were unique to growth on n-hexadecane, and YALI0F23793p was the only protein expressed when grown on glucose (Figure 6M). Also included in Figure 6M was the only annotated fatty acid-CoA ligase (YALI0D17864p) that would convert fatty acids to fatty acyl-CoA. This protein is present for both growth on glucose and n-hexadecane (Figure 6M). *Y. lipolytica* utilized 7 out of 8 proteins in the beta-oxidation pathway where three of these proteins, including YALI0E27654p, YALI0C23859p, and YALI0E06567p were unique to growth on n-hexadecane.

In exploring the other KEGG pathways, we observed a similar trend for propionate (C3) metabolism (Figures 6D, S5), which is unexpected for growth on n-hexadecane that contains an even and saturated carbon chain. For the Krebs or citrate cycle, which is important for energy generation, only HP24 cells, but not HB24 cells, allocated more proteome resources than GP24 cells by 1.4-fold (Figure 6E). Both HP24 and HB24 cells demanded more proteome allocation in the glyoxylate pathway than GP24 cells (Figure 6F), which is known to be important for growth on hydrophobic substrates (e.g., hydrocarbons, lipids, fatty acids) (56) and C2 metabolism (i.e., acetate) (65). In contrast, pyruvate metabolism, which is responsible for converting pyruvate to acetyl CoA, exhibited reduced proteomic resource demand when cells grew on n-hexadecane (Figure 6G). The combined upregulation of the Krebs cycle and downregulation of pyruvate metabolism helps explain the observed high citrate production when cells grew on n-hexadecane (Figure 3B). Proteome allocation for glycolysis/gluconeogenesis and pentose phosphate pathway remained similar regardless of cells growing on either hexadecane or glucose (Figures 6H, 6I).

#### Growth inhibition of *Y. lipolytica* by DP oil can be explained by proteome reallocation

To better understand the inhibitory effect of DP oil, we compared the proteomes of oil-bound HB24 and OB48 cells growing on n-hexadecane and DP oil, respectively (Figures 7A-B). At a high level, we observed distinct proteomes across HB24 and OB48 cells as depicted by the PCA plot (Figure 7C) and differential expression of a large number of proteins (Figures 7D, Data Set S4). Pairwise comparison between HB24 and OB48 cells showed that the mass fraction of amino acids in the measured proteome of OB48 cells increased by 5% or more for Met (+5.48%), His (+6.80%), and Cys (+14.09%) relative to HB24 (Figure 7E).

**Figure 7:**
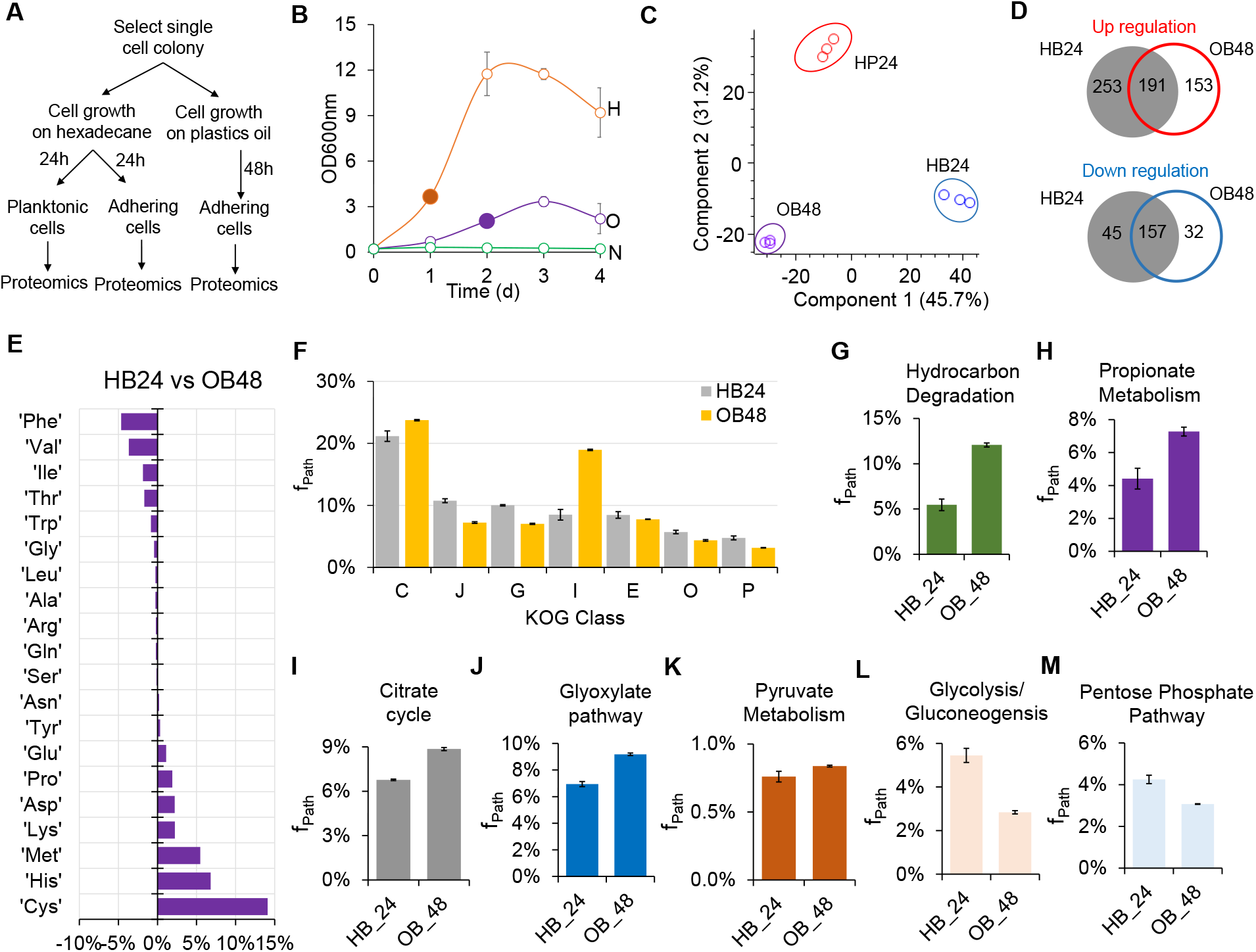
Imbalanced proteome reallocation inhibiting growth of *Y. lipolytica* on DP oil. **(A)** Experimental design. **(B)** Growth and cell sample collection for proteomic analysis. The filled circle indicates that samples were collected for proteomic analysis. Abbreviations: H, hexadecane; O, DP oil; N, no carbon. **(C)** Principal component analysis of proteomes of HP24, HB24, and OB24 cells. **(D)** Proteome comparison between HP24, HB24, and OB48 cells. Regulated proteins between two biological conditions are defined with p-values less than 0.05 and log_2_ fold changes that are either greater than 1 (upregulated proteins) or less than −1 (downregulated proteins). **(E)** Percent change of mass fraction of amino acids in the measured proteomes for comparison between HB24 and OB48. **(F-N)** Comparison of mass fractions of proteomes invested in **(F)** the top 7 KOG classes, **(G)** hydrocarbon degradation pathway, (**H**) propionate metabolism, **(I)** citrate cycle, **(J)** glyoxylate pathway, **(K)** pyruvate metabolism, **(L)** glycolysis/gluconeogenesis, and **(M)** pentose phosphate pathway for growth on n-hexadecane and DP oil.

In contrast to growth on glucose and n-hexadecane, the energy production and conversion KOG class C (f_Path_ = 23.74%) and the lipid transport and metabolism KOG class I (f_Path_ = 18.95%) exhibited the highest proteome allocations among the top 7 KOG classes (C, J, G, I, E, O, and P) that represent 70-75% of the total measured proteomes (Figure 7F). For growth on DP oil that contains a complex mixture of hydrocarbons, we observed a significant proteome reallocation towards the KOG classes C and I by 12% and 123%, respectively at the expense of other KOG classes (J, G, E, O, and P) that are important for growth and cellular biosynthesis. For instance, translation, ribosomal structure and biogenesis (KOG class J), posttranslational modification, protein turnover, chaperones (KOG class O), and amino acid transport and metabolism (KOG class E) showed decreases in proteome allocation by 33%, 24%, and 8%, respectively. We also observed that KOG classes G (carbohydrate transport and metabolism) and P (inorganic ion transport and metabolism) were significantly reduced by 30% and 33%, respectively. Taken together, the rebalanced resource investment imparted by an over-investment in KOG classes C and I might have contributed to the growth inhibition observed when *Y. lipolytica* grew on DP oil (Figure 7B).

Further examination of the metabolism of the KOG classes C and I show that OB48 cells exhibited significant proteome reallocation towards both the hydrocarbon degradation pathway and propionate metabolism by 2.21-fold (p-value = 0, Student’s T-test) and 1.65-fold (p-value = 0.001) respectively, as compared to the HB24 cells (Figures 7G, 7H). The Krebs cycle, glyoxylate shunt, and pyruvate metabolism were also increased by 1.31-fold (p-value = 0), 1.32-fold (p-value = 0), and 1.10-fold (p-value = 0.0137), respectively (Figures 7I, 7J, 7K). In contrast, both glycolysis/gluconeogenesis and pentose phosphate pathway were reduced by 1.92-fold (p-value = 0.0001) and 1.38 (p-value = 0.0003), respectively (Figure 7L, 7M). These results further explained the negative effect of a complex mixture of DP oil on the growth of *Y. lipolytica*.

## DISCUSSION

Increased demand and production of plastic has led to a significant increase in plastic waste. The negative environmental impact of plastic waste necessitates sustainable and economical strategies to recycle it. This study aimed to elucidate and harness robust metabolic capabilities of *Y. lipolytica* for biological upcycling of polyethylene derived from plastic waste into high-value chemicals. To overcome slow kinetics of polyethylene degradation by microorganisms, we demonstrated that hybrid chemical and biological catalysis can offer a promising route for polyolefins upcycling, especially by improving the rate of polymer deconstruction from years to days. Using SiC as a catalyst, catalytic depolymerization of polyethylene resulted in oil/wax suitable for biological conversion. By maintaining DP oil in frozen storage conditions to avoid oil degradation and adapting *Y. lipolytica* to DP oil, we demonstrated that *Y. lipolytica* could grow and produce citric acid and neutral lipids from DP oil. *Y. lipolytica* could effectively consume a wide range of saturated and unsaturated hydrocarbons in DP oil/wax, some of which have not yet been characterized. By analyzing and comparing proteomes of cells growing on glucose, n-hexadecane, and DP oil, we elucidated the unique metabolic capabilities of *Y. lipolytica* for upcycling polyolefins into high-value chemicals.

To utilize hydrocarbons, *Y. lipolytica* grew as a mixture of oil-bound and planktonic cells. This phenomenon occurred once the hydrophobic layer became saturated with bound cells (51). Consistent with literature (66), we observed direct evidence via microscopic images of a surface-mediated transport mechanism enabling cells to access and assimilate hydrocarbons (Figure 2E). The two cell populations exhibited very distinct proteomes (Figure 5C). We found that proteome reallocation towards both energy and lipid metabolism was critical for robust growth of *Y. lipolytica* on DP oil with n-hexadecane as the most preferential substrate (Figures 5, 6, 7). *Y. lipolytica* expressed and upregulated many different proteins in the upstream catabolic pathways (i.e., the hydrocarbon degradation pathway, Krebs cycle, electron transport chain, and propionate metabolism). For instance, the oxysterol-binding protein (Q6C4Y2, YarlipO1F2_226951, or YALI0E22781p) of lipid metabolism was upregulated by 68-fold, which might have played a significant role in hydrocarbon uptake. Likewise, the mitochondrial F1F0-ATP synthase (Q6C877, YarlipO1F2_232437, YALI0D22022p) of energy metabolism was upregulated by 376-fold, which is critical for energy generation from NADH derived from the beta-oxidation of fatty acids.

While DP oil contains rich carbon and energy sources, it is inhibitory to microorganisms. One source of microbial inhibition was caused by chemical inhibitors generated due to the degradation of DP oil. Adapted *Y. lipolytica* could not grow on DP oil stored at room temperature for a long period of time (> 3 months). The FTIR, NMR, and GCMS analyses suggested that ketones likely inhibited cell growth. Based on literature, formation of ketones at room temperature are possible and occur through radical reactions under ambient UV light between atmospheric oxygen and the unsaturated hydrocarbons (67, 68). However, DP oil was stable and did not inhibit cell growth if stored frozen immediately after catalytic depolymerization.

Another source of microbial inhibition was caused by the complex mixture of hydrocarbons (i.e., alkanes and alkenes) in DP oil. Wildtype *Y. lipolytica* exhibited poor growth on DP oil even with glucose supplementation (Figures 2B), indicating that the toxic components (e.g., shorter chain n-alkanes and alkenes) in DP oil might have inhibited cell growth (Figures 2F, S2). Short-term adaptation improved cell growth on both DP oil and n-hexadecane (a model hydrocarbon substrate) (Figures 2C, 2D), suggesting hydrocarbon metabolism was cryptic and required activation as previously observed in both hydrophobic (69) and hydrophilic (70) substrate metabolism. *Y. lipolytica* grew more robustly on n-hexadecane than DP oil because utilizing DP oil required expression of more specialized enzymes to assimilate the mixture of complex hydrocarbons. Proteome analysis suggested that a significant increase in resource investment of energy and lipid metabolism at the expense of protein synthesis may have resulted in the observed growth inhibition. Tuning metabolic fluxes for both the catabolic and anabolic processes together with culture condition optimization are thus critical for balancing the resource allocation and hence improving the upcycling of DP oil in future studies. One simple but effective strategy is to perform long-term adaptive laboratory evolution of *Y. lipolytica* growing on DP oil based on growth selection.

Albeit with room for improvement, *Y. lipolytica* showed promise as a biomanufacturing platform for upcycling plastic waste into higher-value chemicals such as citric acid and neutral lipids (Figure 4). We observed drastic differences in cell growth and hydrocarbon assimilation between cultures growing on n-hexadecane and DP oil. Cells grown with n-hexadecane consumed all of the substrate, produced ∼6-fold more citric acid, and achieved ∼3.5 greater cell mass as compared to cells grown on DP oil. Interestingly, while GCMS analysis revealed most of the alkanes and alkenes in DP oil were consumed, DP oil-grown cells did not achieve similar cell growth or citric acid titers as compared to n-hexadecane-grown cells. We speculate that assimilated DP oil was either converted and secreted as an extracellular emulsifier (e.g., liposan) (71), functionalized into an intermediate metabolite of alkane/alkene degradation (i.e., aldehydes, alcohols or fatty acids), and/or respired as carbon dioxide. Interestingly, *Y. lipolytica* did not grow on individual short-chain (i.e., C11) and long-chain (i.e., C22) 1-alkenes (Figures 2F, S2), although these were assimilated when present in the more complex DP oil. Therefore, the question arises about why *Y. lipolytica* lacks the metabolic capacity required to support cell growth on these specific alkenes? This represents a major challenge necessary to optimize cell growth and product formation from DP oil, which will be investigated in our future studies.

In summary, our work reveals that *Y. lipolytica* has metabolic capabilities that position it as a biomanufacturing platform for upcycling plastic waste-derived DP oil into higher-value chemicals. Our presented findings will undoubtedly help optimize catalytic depolymerization parameters to target medium-chain hydrocarbons while avoiding the generation of potentially unstable and/or inhibitory components that impact efficient biological upcycling. Future work focusing on the specific mechanisms and genes governing accessibility, uptake, and regulation of hydrocarbon metabolism is critical to unlock the full potential of *Y. lipolytica* for upcycling plastic waste (72). Rewiring robust metabolism of *Y. lipolytica* to achieve balanced resource investment for DP oil upcycling is necessary to overcome growth inhibition through rational strain engineering and/or adaptive laboratory evolution.

## MATERIALS AND METHODS

### Strains

The parent *Y. lipolytica* Po1f was obtained from the American Type Culture Collection (ATCC MYA-2613) and was used as the parent strain for short-term adaptation to create the adapted *Y. lipolytica* strain.

### Media and reaction conditions

#### Catalytic depolymerization of plastic waste

The catalytic cleavage of LLDPE reaction was performed under CO_2_ at atmospheric pressure to generate depolymerized plastic (DP) oil. The catalyst reaction contained LLDPE (cat# PE-NA206000, LyondellBasell, TX, USA) and were mixed with 100 mesh unmodified SiC particles (cat# 93-1432, Strem Chemical, MA, USA) (Figure 1B). LLDPE has an average molecular weight of 155,000 g/mol with 18 to 21 branch points per 1000 carbon atoms where branches are comprised of methyl, ethyl, butyl, amyl, and hexyl groups). The entire reactor was vacuumed with a roughing pump for 30 minutes and then back filled with Bone Dry grade CO_2_ to atmospheric pressure three times. The catalyst bed was then heated to 400°C in 30 minutes and held for 8 hours with the condenser side kept at −77°C during the course of the reaction to promote condensation flow of the products. Following the reaction, the oil and wax were separated and purged with N_2_ and stored at −20°C. In the case of catalyst added reactions, the grease was extracted from SiC using a hot toluene bath.

#### Media for cell culturing

Defined media contained yeast nitrogen base without amino acids (Sigma #Y0626, USA) and supplemented with 380 mg/L leucine and 76 mg/L uracil. Nitrogen-limited media contained yeast nitrogen base without amino acids and ammonium sulfate (Sigma #Y1251, USA) and supplemented with 380 mg/L leucine, 76 mg/L uracil, 100 mM HEPES (Fisher #BP310, USA), 90 mM dibasic phosphate, 10 mM monobasic phosphate, 0.73 g/L ammonium sulfate and adjusted to pH of 5 with 6 N hydrochloric acid. Glucose, DP oil, and individual hydrocarbons (Fisher Scientific, USA) were used as carbon sources at concentrations mentioned throughout the text.

#### Cell culturing conditions

All cell culturing experiments were conducted using a MaxQ6000 air incubator set to 250 rpm and 28 ℃. Seed cultures were generated by inoculating a single colony from a fresh petri dish in 3 mL of defined media of a 14 mL culture tube containing 20 g/L glucose and incubated overnight. Sub-seed cultures were generated by transferring 1.5 mL of the seed cultures into 50 mL of defined media of a 500 mL baffled flask containing 20 g/L glucose and incubated overnight until the mid-exponential growth phase. For nitrogen-limited experiments, sub-seed cultures were incubated for 2 days in nitrogen-limited media containing 20 g/L glucose. Cells were centrifuged and washed once in sterile water prior to starting main experiments in 50 mL baffled flasks with a 10 mL working volume and 3 technical replicates. For individual hydrocarbon experiments, 6mM of each hydrocarbon was added to defined media.

#### Short-term adaptation of Y. lipolytica in DP oil

*Y. lipolytica* Po1f was first cultured in defined media with 2% (v/v) DP oil until growth plateaued. Cells were centrifuged at 3500 rpm for 3 minutes and resuspended at 0.25 OD600_nm_ in a new flask containing fresh defined media with 2% (v/v) DP oil. Cells were incubated until achieving an OD greater than 1, and the top-performing replicate was transferred into a new flask containing defined media with 2% (v/v) DP oil. This transfer was repeated for a total of 5 times to generate a DP oil-adapted *Y. lipolytica* strain which was used for all succeeding experiments.

### Analytical methods

#### Cell growth

All cell growth results were quantified by measuring optical density at 600 nm absorbance (OD_600nm_) of 250 µL cell culture in a 96-well plate using a BioTek Synergy microplate reader.

#### Lipid quantification

A 1.2 mL volume of cell cultures was transferred into a pre-weighted 1.5 mL centrifuge tube, washed twice with water, and resuspended to a total volume of 1.2 mL in water. 100 µL aliquots of each sample were transferred into a single well in duplicate using a 96- well plate for lipid quantification. The remaining 1 mL of the cell culture sample was centrifuged at maximum speed (13,000 x g) for 5 minutes before removing the supernatant to determine dry cell weight (DCW, g/L). The cell pellet was dried at 50 ℃ until a constant weight was obtained (e.g., 2-3 days) and used to calculate dry cell weight (DCW). To determine lipid content, a corn oil standard was prepared at a 5 mg/mL concentration in ethanol and used to generate working standards ranging 0.05-1 mg/mL in water. 2 µL of 1 µg/mL BODIPY 435/503 (Fisher #D3922, USA) in DMSO was added to each well of a microplate containing either cells or oil standard, and the plate was shaken in the dark for 15 minutes before measuring fluorescence (excitation 485nm; emission 528nm). Percent of lipid accumulated inside the cell (% Lipid by weight) was calculated by dividing the lipid concentration (g/L) by DCW (g/L).

#### High Performance Liquid Chromatography (HPLC)

Metabolites (i.e., mono sugars, carboxylic acids) were quantified by HPLC. 1 mL cell culture samples were centrifuged at 13,000 x g for 2 minutes and the supernatant was transferred to a new centrifuge tube. Glucose and organic acids were measured after filtering supernatant samples with 0.2 µm filters using a Shimadzu HPLC system equipped with UV and RID detectors (Shimadzu Scientific Instruments, MD, USA) and an Aminex 87H column (Biorad, CA, USA) at a flow rate of 0.6 mL/min and oven temperature of 48 ℃ with 10 mN sulfuric acid mobile phase (73).

#### Gas chromatography coupled with mass spectrometer (GCMS)

Hydrocarbons were quantified by GCMS. 10 mL of sacrificial flask replicates were transferred into a 15 mL tube and stored at – 20 ℃. Samples were thawed to room temperature before adding 2 mL of solvent (i.e., chloroform containing 100 mg/L ethyl pentadecanoate as the internal standard). Samples were vortexed and incubated at room temperature for 2 hours prior to centrifuging at 3500 rpm for 5 minutes, extracting the organic phase, and filtering through a 0.2 μm filter. For samples cultured with n-hexadecane as the sole carbon source, 10 µL of extract was transferred into a GC vial containing 1990 µL of solvent. For samples cultured with DP oil, 125 µL of extract was transferred into a GC vial with an insert containing 125 µL solvent. The GC (HP 7820A, Agilent, CA, USA) equipped with a MS (HP 5977B, Agilent, CA, USA) method was used to detect n-hexadecane and DP substrates as follows: 1 µL of each sample was injected into the GC capillary column (HP- 5MS, 30 m x 0.25 mm x 0.25 µm, Agilent, CA, USA) using spitless mode at injection temperature of 300 °C. The carrier gas, helium, was set at a flow rate of 14.7 mL/min. The oven was set to an initial temperature of 80 °C and held for 5 min, ramped by 30 °C/min to 100 °C and held for 12 mins, ramped by 40 °C /min to 160 °C and held for 15 min, ramped by 40 °C/min to 300 °C and held for 22 min, and baked at 325 °C for 10 min. Peak areas of extracted ions at specific retention times were used to quantify substrates as illustrated in Data Set S1. These peak areas were normalized for each individual sample by dividing by the peak area of the respective internal standard.

#### Liquid chromatography tandem mass spectrometer (LCMS/MS)-based proteomic measurements

*Y. lipolytica* was grown on glucose, hexadecane, or DP oil as described above and sampled at 24 and 48 hours. Cultures were centrifuged at 4700 x g for 10 minutes to isolate distinct populations of cells via phase separation (Figure 2E). Oil-bound cells and culture supernatants were removed from the pelleted planktonic cells, and adherent cells isolated from supernatants via filtration. Planktonic cells and bound cells were then processed for liquid chromatography-tandem mass spectrometry (LC-MS/MS)-based shotgun proteomics as previously detailed (49, 74). Briefly, cells were lysed via bead beating (0.5 mm zirconium oxide beads) in 100 mM Tris-HCl, pH 8.0 (Geno/Grinder 2010; SPEX). Crude lysates were then mixed with 4% SDS and 10 mM dithiothreitol and then heated to 90°C for 10 minutes. Crude lysates were then pre-cleared via centrifugation at 21,000 x g for 10 minutes, moved to a new microfuge tube, adjusted to 30 mM iodoacetamide, and incubated for 20 min at room temperature in the dark. Proteins were isolated to magnetic beads (1 micron SeraMag beads; GE Healthcare) by the protein aggregation capture (PAC) method (75), washed with acetonitrile and ethanol, and digested *in situ* with sequencing-grade trypsin (1:75 (w/w) trypsin to protein ratio) in 100 mM ammonium bicarbonate, pH 8.0 at 37°C overnight. Samples were digested again the following day for 4 h. The tryptic digests were then acidified to 0.5% formic acid, filtered through a 10 kDa MWCO spin filter (Vivaspin500; Sartorius), and quantified via Nanodrop OneC at A205 nm.

Peptide samples were analyzed by automated 1D LC-MS/MS analysis using a Vanquish UHPLC plumbed directly to a nanoelectrospray source installed on a Q Exactive Plus mass spectrometer (Thermo Scientific). Peptide samples were de-salted and separated using a trapping column coupled to an in-house pulled nanospray emitter, as previously described (74). The trapping column (100 µm ID) was packed with 6 cm of 5 µm Kinetex C18 reversed phase (RP) resin (Phenomenex), while the nanospray emitter (75 µm ID) was packed with 15 cm of 1.7 mm Kinetex C18 RP resin. For each sample, 3 µg of peptides were loaded, desalted, and separated by UHPLC along a 180-minute organic gradient (49). Eluting peptides were measured and sequenced by data-dependent acquisition on the Q Exactive MS as previously described (76).

All raw mass spectra for quantification of proteins used in this study have been deposited in the MassIVE and ProteomeXchange data repositories under accession numbers MSV000091396 (MassIVE) and PXD040555 (ProteomeXchange), with data files available at ftp://massive.ucsd.edu/MSV000091396/.

#### Fourier transform infrared (FTIR) spectroscopy

The FTIR spectra were obtained by an ALPHA II Bruker from a thin film of a neat sample from the −20°C and the room temperature stored DP oil over a diamond window. The FTIR spectra corresponds to the sum of 400 scans at a 4 cm^−1^ spectral resolution.

#### Nuclear magnetic resonance (NMR)

NMR experiments were performed at room temperature on a Varian VNMRS 500 MHz spectrometer. Processing of the spectra was performed using the software MNova. For sample preparation, 200 μL of the DP oil were diluted in 400 μL of deuterated chloroform (Sigma-Aldrich 99.8%).

#### Gel permeation chromatography (GPC)

Gel permeation chromatography (GPC) was used to characterize the molecular weight distribution for the grease samples. The tests were carried out using a Viscotek 350A HT-GPC equipped with RI detection and using 1,2,4- trichlorobenzene (TCB) at 145°C as the mobile phase.

### Bioinformatics and *c*omputational analysis

#### Proteomic analysis

MS/MS spectra were searched against the *Y. lipolytica* pO1F v2 proteome (JGI) (77) appended with common protein contaminants using the SEQUEST HT algorithm in Proteome Discoverer v.2.3 (Thermo Scientific). Peptide spectrum matches (PSM) were required to be fully tryptic, allowing for up to 2 miscleavages; a static modification of 57.0214 Da on cysteine (carbamidomethylation) and a dynamic modification of 15.9949 Da on methionine (oxidation) residues. PSMs were scored and filtered using Percolator with false-discovery rates (FDR) initially controlled at <1% at the PSM-and peptide-level. Peptides were then quantified by chromatographic area-under-the-curve, mapped to their respective proteins, and areas summed to estimate protein-level abundance. For proteome reallocation analysis, these protein abundances were used directly for the calculation without filtering and imputation. For differential expression analysis, proteins that do not have at least 70% valid values in each group were removed and remaining proteins were log2 transformed using Perseus v2.0.6.0 (78). Missing values were imputed to estimate the mass spectrometer’s limit of detection in Perseus. Statistical tests (T-test and ANOVA) were performed with default parameter settings in Perseus. Significant differences in protein abundance were calculated for relevant sample group comparisons after FDR correction.

Though the Po1f strain and its associated proteome database were interrogated in this study, more detailed functional information is available for the *Y. lipolytica* CLIB122 reference proteome. To improve functional annotations and to view the results in the same context as the CLIB122 reference strain, the Po1f strain was analyzed with OmicsBox v2.2.4 software. This enhanced search provided BLASTP, InterPro, GO, EC, KEGG, KOG, and Signal P information, and enabled the generation of an orthology lookup table linking Po1f proteins to their orthologs in the CLIB122 reference proteome.

#### Proteome reallocation analysis

KEGG and Uniprot were used to obtain protein annotation, protein sequences, and pathway ontology (79, 80) (81). The hydrocarbon degradation pathway that oxidizes hydrocarbons to fatty acids in endoplasmic reticulum and then acetyl CoA in peroxisome were manually curated based on literature sources (56, 57, 59-61).

Mass fraction of a protein P_i_, f_Pi_, in the proteome is calculated by (82)

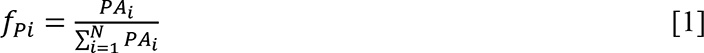

where PA_i_ is the abundance of protein i, N is the total number of proteins in the measured proteome, and 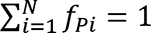. Mass fraction of a pathway proteome, f_Path_, that contains M proteins is determined by (82):

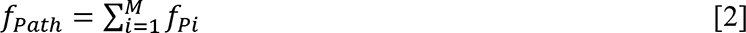

Mass fraction of amino acid j, f_Aj_, in the proteome is computed by (82):

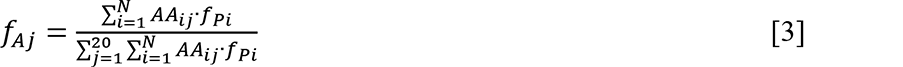

where AA_ij_ is the abundance of amino acid j in protein i of the proteome and 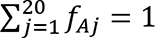.

## ACKNOWLEDGEMENTS

This research is financially supported in part by the ORII Seed fund at The University of Tennessee, Knoxville (to CTT, BK, SL, and RG) and the DOE BER Genomic Science Program (DE-SC0019412 to CTT and RG). The views, opinions, and/or findings contained in this article are those of the authors and should not be interpreted as representing the official views or policies, either expressed or implied, of the funding agencies. Mention of trade names or commercial products in this publication is solely for the purpose of providing specific information and does not imply recommendation or endorsement by the funding agencies.

## SUPPLEMENTARY MATERIALS

**Figure S1:** NMR spectra for fresh and 3-month DP oil/wax samples analyzed by 1H-13C HMBC **(A)** Full NMR spectrum. **(B)** Zoom-in spectrum for 1H 0-3ppm and 13C 0-60ppm ranges.

**Figure S2:** Growth kinetics of adapted *Y. lipolytica* on (**A-P**) n-alkanes and (**Q-T**) 1-alkenes. The growth experiments were performed with at least 6 biological replicates. Each value on a graph is represented by an average ± 1 standard deviation (n = 6 or 12).

**Figure S3:** Comparison of proteomes of *Y. lipolytica* growing on glucose and n-hexadecane. **(A-C)** Volcano plots depict differential protein expression between **(A)** GP24 and HP24 cells, **(B)** GP24 and HB24 cells, and **(C)** HP24 and HB24 cells. **(D)** Heatmap showing distinct clusters of proteins that were differentially expressed for growth on different substrates.

**Figure S4:** Comparison of mass fractions of proteomes invested in all 23 KOG classes of *Y. lipolytica* growing on glucose, n-hexadecane, and DP oil.

**Figure S5:** Proteome reallocation for propionate metabolism of *Y. lipolytica* growing on glucose, n-hexadecane, and DP oil. **(A)** Metabolic pathway of propionate degradation. **(B)** Mass fractions of proteins involved in the propionate metabolism.

**Data Set S1:** Pairwise comparison of differential protein expression between the planktonic GP24 and planktonic HP24 cells.

**Data Set S2:** Pairwise comparison of differential protein expression between the planktonic GP24 and bound HB24 cells.

**Data Set S3:** Pairwise comparison of differential protein expression between the planktonic HP24 and bound HB24 cells.

**Data Set S4:** Pairwise comparison of differential protein expression between the bound HB24 and bound OB24 cells.

## REFERENCES

1. Geyer R, Jambeck JR, Law KL. 2017. Production, use, and fate of all plastics ever made. Science advances 3:e1700782.

2. Law KL, Starr N, Siegler TR, Jambeck JR, Mallos NJ, Leonard GH. 2020. The United States’ contribution of plastic waste to land and ocean. Science Advances 6:eabd0288.

3. Beydoun K, Klankermayer J. 2020. Efficient Plastic Waste Recycling to Value-Added Products by Integrated Biomass Processing. ChemSusChem 13:488–492.

4. Ellis LD, Rorrer NA, Sullivan KP, Otto M, McGeehan JE, Román-Leshkov Y, Wierckx N, Beckham GT. 2021. Chemical and biological catalysis for plastics recycling and upcycling. Nature Catalysis 4:539–556.

5. Sullivan KP, Werner AZ, Ramirez KJ, Ellis LD, Bussard JR, Black BA, Brandner DG, Bratti F, Buss BL, Dong X, Haugen SJ, Ingraham MA, Konev MO, Michener WE, Miscall J, Pardo I, Woodworth SP, Guss AM, Román-Leshkov Y, Stahl SS, Beckham GT. 2022. Mixed plastics waste valorization through tandem chemical oxidation and biological funneling. Science 378:207–211.

6. Gambarini V, Pantos O, Kingsbury JM, Weaver L, Handley KM, Lear G. 2021. Phylogenetic Distribution of Plastic-Degrading Microorganisms. mSystems 6:e01112–20.

7. Danso D, Chow J, Streit WR. 2019. Plastics: Environmental and Biotechnological Perspectives on Microbial Degradation. Applied and Environmental Microbiology 85:e01095–19.

8. Yoshida S, Hiraga K, Takehana T, Taniguchi I, Yamaji H, Maeda Y, Toyohara K, Miyamoto K, Kimura Y, Oda K. 2016. A bacterium that degrades and assimilates poly (ethylene terephthalate). Science 351:1196–1199.

9. Chamas A, Moon H, Zheng J, Qiu Y, Tabassum T, Jang JH, Abu-Omar M, Scott SL, Suh S. 2020. Degradation rates of plastics in the environment. ACS Sustainable Chemistry & Engineering 8:3494–3511.

10. Ellis LD, Orski SV, Kenlaw GA, Norman AG, Beers KL, Román-Leshkov Y, Beckham GT. 2021. Tandem Heterogeneous Catalysis for Polyethylene Depolymerization via an Olefin-Intermediate Process. ACS Sustainable Chemistry & Engineering 9:623–628.

11. Lynd LR, Beckham GT, Guss AM, Jayakody LN, Karp EM, Maranas C, McCormick RL, Amador-Noguez D, Bomble YJ, Davison BH. 2022. Toward low-cost biological and hybrid biological/catalytic conversion of cellulosic biomass to fuels. Energy & Environmental Science 15:938–990.

12. Beltrame PL, Carniti P, Audisio G, Bertini F. 1989. Catalytic degradation of polymers: Part II—Degradation of polyethylene. Polymer Degradation and Stability 26:209–220.

13. Ding W, Liang J, Anderson LL. 1997. Thermal and catalytic degradation of high density polyethylene and commingled post-consumer plastic waste. Fuel Processing Technology 51:47–62.

14. Karagöz S, Yanik J, Uçar S, Saglam M, Song C. 2003. Catalytic and thermal degradation of high-density polyethylene in vacuum gas oil over non-acidic and acidic catalysts. Applied Catalysis A: General 242:51–62.

15. Keane MA. 2009. Catalytic Transformation of Waste Polymers to Fuel Oil. ChemSusChem 2:207–214.

16. Audisio G, Silvani A, Beltrame PL, Carniti P. 1984. Catalytic thermal degradation of polymers: Degradation of polypropylene. Journal of Analytical and Applied Pyrolysis 7:83–90.

17. Holm MS, Taarning E, Egeblad K, Christensen CH. 2011. Catalysis with hierarchical zeolites. Catalysis Today 168:3–16.

18. Hwang E-Y, Kim J-R, Choi J-K, Woo H-C, Park D-W. 2002. Performance of acid treated natural zeolites in catalytic degradation of polypropylene. Journal of Analytical and Applied Pyrolysis 62:351–364.

19. Corma A. 1995. Inorganic Solid Acids and Their Use in Acid-Catalyzed Hydrocarbon Reactions. Chemical Reviews 95:559–614.

20. Jentoft FC, Gates BC. 1997. Solid-acid-catalyzed alkane cracking mechanisms: evidence from reactions of small probe molecules. Topics in Catalysis 4:1–13.

21. Knaeble W, Carr RT, Iglesia E. 2014. Mechanistic interpretation of the effects of acid strength on alkane isomerization turnover rates and selectivity. Journal of Catalysis 319:283–296.

22. Lukyanov DB, Shtral VI, Khadzhiev SN. 1994. A kinetic model for the hexane cracking reaction over H-ZSM-5. Journal of Catalysis 146:87–92.

23. Macht J, Carr RT, Iglesia E. 2009. Consequences of Acid Strength for Isomerization and Elimination Catalysis on Solid Acids. Journal of the American Chemical Society 131:6554–6565.

24. Macht J, Janik MJ, Neurock M, Iglesia E. 2008. Mechanistic Consequences of Composition in Acid Catalysis by Polyoxometalate Keggin Clusters. Journal of the American Chemical Society 130:10369–10379.

25. Borges P, Pinto RR, Lemos M, Lemos F, Védrine J, Derouane E, Ribeiro FR. 2005. Activity– acidity relationship for alkane cracking over zeolites: n-hexane cracking over HZSM-5. Journal of Molecular Catalysis A: Chemical 229:127–135.

26. Song Y, He Y, Laursen S. 2020. Controlling Selectivity and Stability in the Hydrocarbon Wet-Reforming Reaction Using Well-Defined Ni + Ga Intermetallic Compound Catalysts. ACS Catalysis 10:8968–8980.

27. Song Y, Laursen S. 2019. Control of surface reactivity towards unsaturated CC bonds and H over Ni-based intermetallic compounds in semi-hydrogenation of acetylene. Journal of Catalysis 372:151–162.

28. He Y, Song Y, Laursen S. 2019. The Origin of the Special Surface and Catalytic Chemistry of Ga-Rich Ni3Ga in the Direct Dehydrogenation of Ethane. ACS Catalysis 9:10464–10468.

29. He Y, Song Y, Cullen DA, Laursen S. 2018. Selective and Stable Non-Noble-Metal Intermetallic Compound Catalyst for the Direct Dehydrogenation of Propane to Propylene. Journal of the American Chemical Society 140:14010–14014.

30. Groenewald M, Boekhout T, Neuvéglise C, Gaillardin C, van Dijck PWM, Wyss M. 2014. Yarrowia lipolytica: Safety assessment of an oleaginous yeast with a great industrial potential. Critical Reviews in Microbiology 40:187–206.

31. Fickers P, Benetti PH, Waché Y, Marty A, Mauersberger S, Smit MS, Nicaud JM. 2005. Hydrophobic substrate utilisation by the yeast Yarrowia lipolytica, and its potential applications. FEMS Yeast Research 5:527–543.

32. Gao C, Yang X, Wang H, Rivero CP, Li C, Cui Z, Qi Q, Lin CSK. 2016. Robust succinic acid production from crude glycerol using engineered Yarrowia lipolytica. Biotechnology for biofuels 9:1–11.

33. Kamzolova SV, Chiglintseva MN, Lunina JN, Morgunov IG. 2012. α-Ketoglutaric acid production by Yarrowia lipolytica and its regulation. Applied Microbiology and Biotechnology 96:783–791.

34. Rymowicz W, Fatykhova AR, Kamzolova SV, Rywińska A, Morgunov IG. 2010. Citric acid production from glycerol-containing waste of biodiesel industry by Yarrowia lipolytica in batch, repeated batch, and cell recycle regimes. Applied microbiology and biotechnology 87:971–979.

35. Rywińska A, Rymowicz W. 2010. High-yield production of citric acid by Yarrowia lipolytica on glycerol in repeated-batch bioreactors. Journal of Industrial Microbiology and Biotechnology 37:431–435.

36. Rymowicz W, Rywińska A, Marcinkiewicz M. 2009. High-yield production of erythritol from raw glycerol in fed-batch cultures of Yarrowia lipolytica. Biotechnology Letters 31:377–380.

37. Blazeck J, Hill A, Liu L, Knight R, Miller J, Pan A, Otoupal P, Alper HS. 2014. Harnessing Yarrowia lipolytica lipogenesis to create a platform for lipid and biofuel production. Nature communications 5.

38. Dulermo T, Nicaud J-M. 2011. Involvement of the G3P shuttle and β-oxidation pathway in the control of TAG synthesis and lipid accumulation in Yarrowia lipolytica. Metabolic engineering 13:482–491.

39. Abghari A, Chen S. 2014. Yarrowia lipolytica as an oleaginous cell factory platform for the production of fatty acid-based biofuel and bioproducts. Bioenergy and Biofuels 2:21.

40. Xue Z, Sharpe PL, Hong S-P, Yadav NS, Xie D, Short DR, Damude HG, Rupert RA, Seip JE, Wang J, Pollak DW, Bostick MW, Bosak MD, Macool DJ, Hollerbach DH, Zhang H, Arcilla DM, Bledsoe SA, Croker K, McCord EF, Tyreus BD, Jackson EN, Zhu Q. 2013. Production of omega-3 eicosapentaenoic acid by metabolic engineering of Yarrowia lipolytica. Nat Biotech 31:734–740.

41. Abdel-Mawgoud AM, Markham KA, Palmer CM, Liu N, Stephanopoulos G, Alper HS. 2018. Metabolic engineering in the host Yarrowia lipolytica. Metabolic engineering 50:192–208.

42. Markham KA, Alper HS. 2018. Synthetic Biology Expands the Industrial Potential of Yarrowia lipolytica. Trends in Biotechnology 36:1085–1095.

43. Epova EG, Marina; Kovalyov, Leonid; Isakova, Elena; Deryabina, Yulia; Belyakova, Alla; Zylkova, Marina and Alexei Shevelev. 2012. Identification of Proteins Involved in pH Adaptation in Extremophile Yeast Yarrowia lipolytica, Proteomic Applications in Biology. InTech.

44. Andreishcheva E, Isakova E, Sidorov N, Abramova N, Ushakova N, Shaposhnikov G, Soares M, Zvyagilskaya R. 1999. Adaptation to salt stress in a salt-tolerant strain of the yeast Yarrowia lipolytica. Biochemistry c/c of Biokhomiia 64:1061–1067.

45. Walker C, Ryu S, Trinh CT. 2019. Exceptional solvent tolerance in Yarrowia lipolytica is enhanced by sterols. Metabolic engineering 54:83–95.

46. Ryu S, Labbé N, Trinh CT. 2015. Simultaneous saccharification and fermentation of cellulose in ionic liquid for efficient production of α-ketoglutaric acid by Yarrowia lipolytica. Applied microbiology and biotechnology 99:4237–4244.

47. Walker C, Ryu S, Garcia S, Dooley D, Mendoza B, Trinh CT. 2022. Gene Coexpression Connectivity Predicts Gene Targets Underlying High Ionic-Liquid Tolerance in Yarrowia lipolytica. mSystems 7:e00348–22.

48. Quarterman J, Slininger PJ, Kurtzman CP, Thompson SR, Dien BS. 2016. A survey of yeast from the Yarrowia clade for lipid production in dilute acid pretreated lignocellulosic biomass hydrolysate. Applied Microbiology and Biotechnology:1–16.

49. Walker C, Dien B, Giannone RJ, Slininger P, Thompson SR, Trinh CT, Tullman-Ercek D. 2021. Exploring Proteomes of Robust Yarrowia lipolytica Isolates Cultivated in Biomass Hydrolysate Reveals Key Processes Impacting Mixed Sugar Utilization, Lipid Accumulation, and Degradation. mSystems 6:e00443–21.

50. Wolf K, Barth G, Gaillardin C. 1996. Yarrowia lipolytica. Springer.

51. Thevenieau F, Beopoulos A, Desfougeres T, Sabirova J, Albertin K, Zinjarde S, Nicaud J-M. 2010. Uptake and assimilation of hydrophobic substrates by the oleaginous yeast Yarrowia lipolytica, Handbook of hydrocarbon and lipid microbiology.

52. Thevenieau F, Le Dall MT, Nthangeni B, Mauersberger S, Marchal R, Nicaud JM. 2007. Characterization of Yarrowia lipolytica mutants affected in hydrophobic substrate utilization. Fungal Genetics and Biology 44:531–542.

53. Mlíčková K, Roux E, Athenstaedt K, d’Andrea S, Daum G, Chardot T, Nicaud J-M. 2004. Lipid accumulation, lipid body formation, and acyl coenzyme A oxidases of the yeast Yarrowia lipolytica. Appl Environ Microbiol 70:3918-3924.

54. Cirigliano MC, Carman GM. 1984. Isolation of a bioemulsifier from Candida lipolytica. Appl Environ Microbiol 48:747–750.

55. Cirigliano MC, Carman GM. 1985. Purification and characterization of liposan, a bioemulsifier from Candida lipolytica. Applied and environmental microbiology 50:846–850.

56. Fickers P, Benetti P-H, Wache Y, Marty A, Mauersberger S, Smit M, Nicaud J-M. 2005. Hydrophobic substrate utilisation by the yeast Yarrowia lipolytica, and its potential applications. FEMS yeast research 5:527–543.

57. Iwama R, Kobayashi S, Ishimaru C, Ohta A, Horiuchi H, Fukuda R. 2016. Functional roles and substrate specificities of twelve cytochromes P450 belonging to CYP52 family in n-alkane assimilating yeast Yarrowia lipolytica. Fungal Genetics and Biology 91:43–54.

58. Iida T, Sumita T, Ohta A, Takagi M. 2000. The cytochrome P450ALK multigene family of an n-alkane-assimilating yeast, Yarrowia lipolytica: cloning and characterization of genes coding for new CYP52 family members. Yeast 16:1077-1087.

59. Gatter M, Förster A, Bär K, Winter M, Otto C, Petzsch P, Ježková M, Bahr K, Pfeiffer M, Matthäus F, Barth G. 2014. A newly identified fatty alcohol oxidase gene is mainly responsible for the oxidation of long-chain ω-hydroxy fatty acids in Yarrowia lipolytica. FEMS Yeast Research 14:858–872.

60. Iwama R, Kobayashi S, Ohta A, Horiuchi H, Fukuda R. 2014. Fatty aldehyde dehydrogenase multigene family involved in the assimilation of n-alkanes in Yarrowia lipolytica. Journal of Biological Chemistry 289:33275–33286.

61. Dulermo R, Gamboa-Meléndez H, Ledesma-Amaro R, Thévenieau F, Nicaud J-M. 2015. Unraveling fatty acid transport and activation mechanisms in Yarrowia lipolytica. Biochimica et Biophysica Acta (BBA)-Molecular and Cell Biology of Lipids 1851:1202–1217.

62. Yang M-L, Zhu Y-A, Fan C, Sui Z-J, Chen D, Zhou X-G. 2011. DFT study of propane dehydrogenation on Pt catalyst: effects of step sites. Physical Chemistry Chemical Physics 13:3257–3267.

63. Saelee T, Namuangruk S, Kungwan N, Junkaew A. 2018. Theoretical Insight into Catalytic Propane Dehydrogenation on Ni(111). The Journal of Physical Chemistry C 122:14678–14690.

64. Wu C, Wang L, Xiao Z, Li G, Wang L. 2020. Understanding deep dehydrogenation and cracking of n-butane on Ni(111) by a DFT study. Physical Chemistry Chemical Physics 22:724–733.

65. Liu N, Qiao K, Stephanopoulos G. 2016. 13C Metabolic Flux Analysis of acetate conversion to lipids by Yarrowia lipolytica. Metabolic engineering 38:86–97.

66. Kim T-H, Oh Y-S, Kim S-J. 2000. The possible involvement of the cell surface in aliphatic hydrocarbon utilization by an oil-degrading yeast, Yarrowia lipolytica 180. Journal of microbiology and biotechnology 10:333-337.

67. Mallégol J, Gardette J-L, Lemaire J. 1999. Long-term behavior of oil-based varnishes and paints I. Spectroscopic analysis of curing drying oils. Journal of the American Oil Chemists’ Society 76:967–976.

68. Rosado EDaP, John. 2003. Chemical characterization of fresh, used and weathered motor oil via GC/MS, NMR and FTIR techniques. Indiana Academy of Science 112.

69. Beier A, Hahn V, Bornscheuer UT, Schauer F. 2014. Metabolism of alkenes and ketones by Candida maltosa and related yeasts. AMB Express 4:75.

70. Ryu S, Hipp J, Trinh CT. 2016. Activating and Elucidating Metabolism of Complex Sugars in <span class="named-content genus-species" id="named-content-1">Yarrowia lipolytica</span>. Applied and Environmental Microbiology 82:1334.

71. Vance-Harrop MH, Gusmão NBd, Campos-Takaki GMd. 2003. New bioemulsifiers produced by Candida lipolytica using D-glucose and babassu oil as carbon sources. Brazilian journal of microbiology 34:120–123.

72. Fukuda R. 2013. Metabolism of hydrophobic carbon sources and regulation of it in n-alkane-assimilating yeast Yarrowia lipolytica. Bioscience, biotechnology, and biochemistry 77:1149–1154.

73. Ryu S, Trinh CT. 2018. Understanding Functional Roles of Native Pentose-Specific Transporters for Activating Dormant Pentose Metabolism in *Yarrowia lipolytica*. Applied and Environmental Microbiology 84:e02146–17.

74. Walker C, Ryu S, Giannone RJ, Garcia S, Trinh CT. 2020. Understanding and Eliminating the Detrimental Effect of Thiamine Deficiency on the Oleaginous Yeast *Yarrowia lipolytica*. Applied and Environmental Microbiology 86:e02299–19.

75. Batth TS, Tollenaere MX, Rüther P, Gonzalez-Franquesa A, Prabhakar BS, Bekker-Jensen S, Deshmukh AS, Olsen JV. 2019. Protein aggregation capture on microparticles enables multipurpose proteomics sample preparation. Molecular & Cellular Proteomics 18:1027–1035.

76. Clarkson SM, Giannone RJ, Kridelbaugh DM, Elkins JG, Guss AM, Michener JK. 2017. Construction and optimization of a heterologous pathway for protocatechuate catabolism in Escherichia coli enables bioconversion of model aromatic compounds. Applied and environmental microbiology 83.

77. Walker C, Ryu S, Haridas S, Na H, Zane M, LaButti K, Barry K, Grigoriev IV, Trinh CT. 2020. Draft Genome Assemblies of Ionic Liquid-Resistant Yarrowia lipolytica PO1f and Its Superior Evolved Strain, YlCW001. Microbiology Resource Announcements 9.

78. Tyanova S, Temu T, Sinitcyn P, Carlson A, Hein MY, Geiger T, Mann M, Cox J. 2016. The Perseus computational platform for comprehensive analysis of (prote) omics data. Nature methods 13:731.

79. Kanehisa M, Goto S. 2000. KEGG: Kyoto Encyclopedia of Genes and Genomes. Nucleic Acids Res 28:27–30.

80. Polpitiya AD, Qian W-J, Jaitly N, Petyuk VA, Adkins JN, Camp DG, II, Anderson GA, Smith RD. 2008. DAnTE: a statistical tool for quantitative analysis of -omics data. Bioinformatics 24:1556-1558.

81. Consortium TU. 2022. UniProt: the Universal Protein Knowledgebase in 2023. Nucleic Acids Research 51:D523–D531.

82. Seo H, Giannone RJ, Yang Y-H, Trinh CT. 2022. Proteome reallocation enables the selective de novo biosynthesis of non-linear, branched-chain acetate esters. Metabolic Engineering.

